# Mismatch resolution for repeat-mediated deletions show a polarity that is mediated by MLH1

**DOI:** 10.1101/2022.07.11.499639

**Authors:** Hannah Trost, Arianna Merkell, Felicia Wednesday Lopezcolorado, Jeremy M. Stark

## Abstract

Repeat-mediated deletions (RMDs) are a type of chromosomal rearrangement between two homologous sequences that causes loss of the sequence between the repeats, along with one of the repeats. Sequence divergence between repeats suppresses RMDs; the mechanisms of such suppression and of resolution of the mismatched bases remains poorly understood. We identified RMD regulators using a set of reporter assays in mouse cells that test two key parameters: repeat sequence divergence and the distances between one repeat and the initiating chromosomal double-strand break. We found that the mismatch repair factor MLH1 suppresses RMDs with sequence divergence in a manner that is epistatic with the mismatch repair factors MSH2 and MSH6, and which is dependent on residues in MLH1 and its binding partner PMS2 that are important for nuclease activity. Additionally, we found that resolution of mismatches in the RMD product have a specific polarity, where mismatched bases that are proximal to the chromosomal break end are preferentially removed. Moreover, we found that the MLH1 endonuclease domain is important for this polarity of mismatch resolution. Finally, we identified distinctions between MLH1 vs. TOP3α in regulation of RMDs. We suggest that MLH1 suppresses RMDs with sequence divergence, while also promoting directional mismatch resolution.

## INTRODUCTION

Repeat-mediated deletions (RMDs) are a type of chromosomal rearrangement involving recombination between two repeat elements that causes a deletion between the repeats, along with one of the repeats (Mendez-Dorantes et al. 2018; Morales et al. 2021). A likely mechanism of RMDs is single-strand annealing (SSA), which involves a chromosomal break between two repeat elements that is resected to generate 3’ ssDNA that enables the two repeat elements to anneal together to bridge the DSB. Subsequent removal of 3’ non-homologous tails, fill-in synthesis, and ligation completes these events (Bhargava et al. 2016).

RMDs have the potential to reshape mammalian genomes, due to the high density of repetitive DNA elements, such as long interspersed elements and short interspersed elements, including approximately one million Alu-like elements in the human genome (Batzer and Deininger 2002; Sen et al. 2006; Han et al. 2007; de Koning et al. 2011). Indeed, RMDs have been associated with several genetic diseases, including loss of tumor suppressor genes leading to increased cancer incidence (Belancio et al. 2010; Burns 2022). Notably, repeat elements show substantial sequence divergence, which is a potent suppressor of recombination between repeat sequences (Waldman and Liskay 1988; Mezard et al. 1992). For example, Alu-like elements can show up to 20% sequence divergence between elements (Batzer and Deininger 2002).

Components of mismatch repair appear to play a key role in suppressing recombination between divergent sequences (Rayssiguier et al. 1989; Alani et al. 1994). The mismatch repair pathway is a critical aspect of DNA replication to excise misincorporated bases (Modrich 2006; Hombauer et al. 2011). This pathway involves recognition of the mismatch by MSH2 in complex with MSH3 or MSH6, which recruits MLH1 in complex with one of its binding partners (e.g., PMS2) to generate a DNA nick upstream of the mismatch, which is generally followed by excision of the nicked strand by EXO1 and/or displacement synthesis (Modrich 2006; Hombauer et al. 2011).

There are apparent mechanistic distinctions between mismatch repair during DNA replication vs. suppression of recombination between divergent sequences. For example in *Saccharomyces cerevisiae*, both *MSH6* and *MLH1* are required for mismatch repair; but only *MSH6* is required for suppression of recombination between divergent sequences, whereas *MLH1* appears dispensable (Sugawara et al. 2004; Goldfarb and Alani 2005; Chakraborty and Alani 2016). Whether these mechanistic distinctions are conserved in mammalian cells has been unclear, as are other aspects of the role of mismatch repair in regulation of RMDs. For example, the mechanisms and patterns of mismatch resolution during RMDs have been poorly understood. Additionally, the relationship between mismatch repair and other factors important for suppression of recombination between divergent sequences is unclear. In this study, we have used an assay system for RMDs in mouse cells to survey the influence of several DNA damage response factors on distinct RMD events, and subsequently focus on defining the role of MLH1 on regulation of RMDs between divergent repeats, including for mismatch resolution.

## RESULTS

### Components of mismatch repair and the BLM-TOP3α-RMI1/2 (BTR) complex suppress RMDs, whereas several other factors promote these events

We sought to identify DNA damage response factors that influence the formation of RMDs, using a reporter system that uses GFP expression as a measure of RMDs, called RMD-GFP (Figure 1A) (Mendez-Dorantes et al. 2018). This reporter has two tandem 287 bp repeats (shown as “R”) separated by 0.4 Mbp on chromosome 17 in mouse embryonic stem cells (mESCs). The 5’ repeat is the endogenous sequence located just downstream of the *Cdkn1A* promoter, and the 3’ repeat is targeted to the *Pim1* locus and is fused to GFP. An RMD between these two repeats generates a *Cdkn1A-GFP* fusion gene that causes GFP+ cells, which can be measured with flow cytometry. To induce an RMD, we introduce two DSBs between the two repeats using Cas9/sgRNAs. The 5’ DSB is always at the same position, which is 268 bp downstream of the 5’ repeat (5’ 268 bp). The 3’ DSB can be made at various distances upstream of the 3’ repeat, which we refer to as the DSB/repeat distance. There are also two other versions of the RMD-GFP reporter that contain equally spaced mismatches in the 3’ repeat: 1%RMD-GFP with 3 mismatches causing 1% sequence divergence, and 3%RMD-GFP with 8 mismatches causing 3% sequence divergence. All assay conditions are normalized to transfection efficiency with parallel transfections with a GFP expression vector.

**Figure 1.**
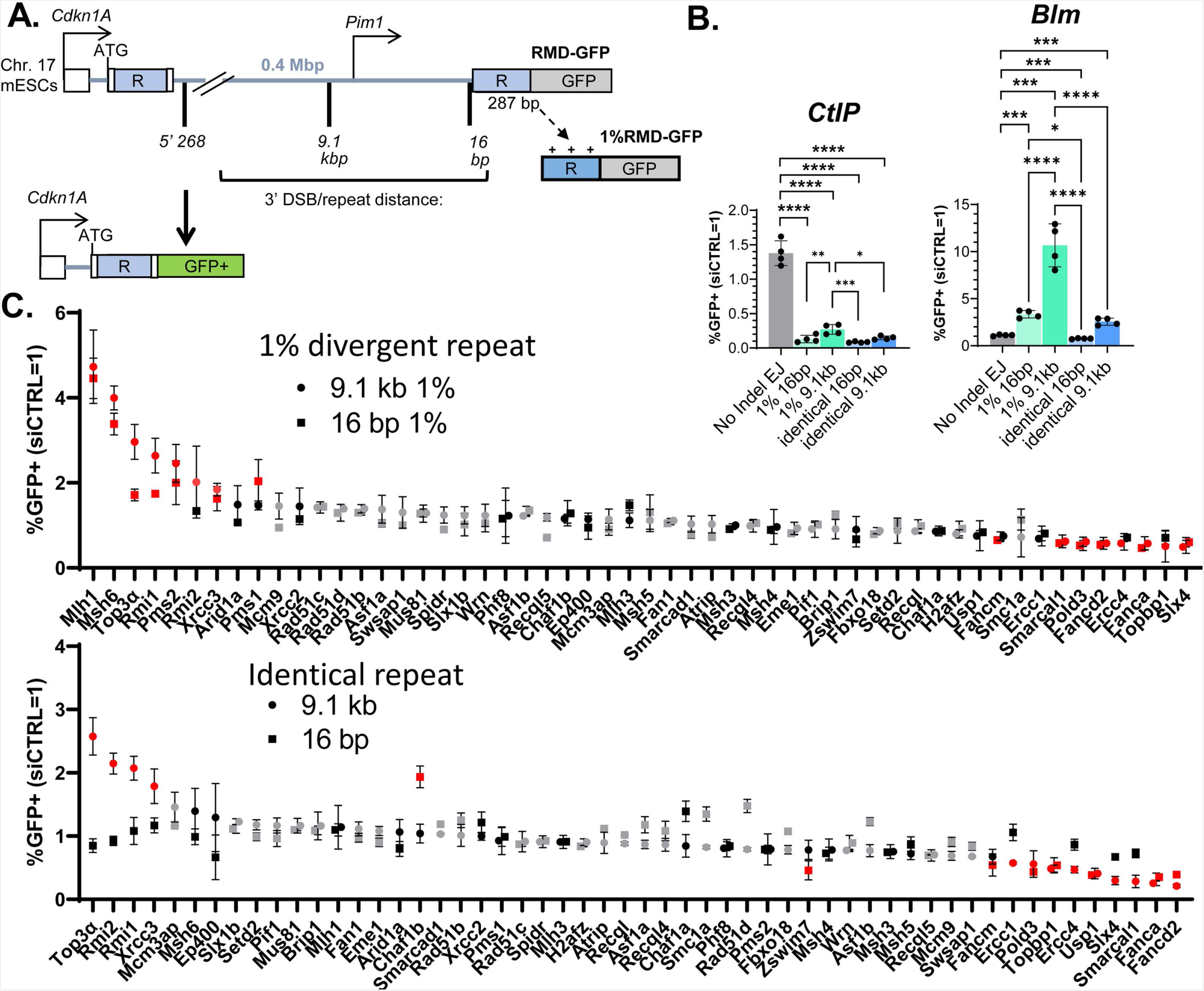
Components of mismatch repair and the BLM-TOP3α-RMI1/2 (BTR) complex suppress RMDs, whereas several other factors promote these events. **(A)** Shown is the RMD-GFP reporter, which is integrated into the *Pim1* locus in chromosome 17 of mESCs, such that repair of two DSBs by an RMD leads to GFP+ cells. The two repeats shown as “R”, the 5’ repeat being endogenous sequence and the 3’ repeat is fused to GFP. 1%RMD-GFP has 1% sequence divergence between the repeats. RMDs are induced by creating two DSBs: one 268 bp downstream of the 5’ repeat, and the other either 16 bp or 9.1 kb upstream of the 3’ repeat, which we refer to as the DSB / repeat distance. **(B)** Shown are the effects of siRNAs targeting BLM (siBlm) and CtIP (siCtIP) for 4 RMD reporter assays, and an NHEJ assay (No Indel EJ). Repair frequencies are normalized to transfection efficiency, and parallel non-targeting siRNA (siCTRL = 1). n=4. *p ≤ 0.05, **p ≤ 0.005, ***p ≤ 0.0005, ****p < 0.0001, Statistics are one-way ANOVA using Tukey’s multiple comparisons test. **(C)** Shown are effects of siRNAs targeting 54 factors on frequency of RMDs at both 16 bp and 9.1 kb DSB/repeat distances for 1%RMD-GFP and RMD-GFP. Frequencies are normalized to transfection efficiency and parallel siCTRL (=1). Genes are ranked by the fold-effect relative to siCTRL at 9.1 kb. All siRNAs tested n=2, and those with ≥1.5-fold effect from these trials were tested a total of n=4. Grey: n=2, Black: n=4, Red: n=4 and also ≥1.5-fold effect relative to siCTRL. Data are represented as mean values ± SD.

To begin with, we examined two factors already implicated in RMD regulation (Mendez-Dorantes et al. 2018; Mendez-Dorantes et al. 2020), which served as controls during our survey of other factors, as described below. Specifically, we examined effects of siRNA depletion of the BLM helicase and the end resection factor CtIP on four versions of the RMD-GFP assay: 1) RMD-GFP with the 16 bp DSB/repeat distance, 2) RMD-GFP with the 9.1 kbp DSB/repeat distance, 3) 1%RMD-GFP with the 16 bp DSB/repeat distance, and 4) 1%RMD-GFP with the 9.1 kbp DSB/repeat distance. We chose these versions of the assay as it enables a comparison of identical vs. divergent repeats, each at both very short and relatively long DSB/repeat distance. We also included an assay for non-homologous end joining (NHEJ) as a control (EJ7-GFP / No Indel EJ assay). With this approach, we found that depleting the end resection factor CtIP causes a significant decrease in all four RMDs compared to NHEJ, although the 1%RMD-GFP (i.e., divergent repeat) assay with the 9.1 kbp DSB/repeat distance was affected the least (Figure 1B). In contrast, BLM depletion caused a specific increase in three of the RMDs (i.e., RMDs with both identical and divergent repeats at the 9.1 kbp DSB/repeat distance, and the divergent repeat at 16 bp), and a modest decrease for the identical repeat at 16 bp, each compared to the effect on NHEJ (Figure 1B). Also, the fold-effects of BLM depletion differed among the RMD events, with the divergent repeat at 9.1 kbp showing a markedly greater effect (Figure 1B). We confirmed siRNA depletion of CtIP and BLM with both qRT-PCR and immunoblotting (Supplemental Figure S1A, S1B).

Using these four variants of the RMD assay, we then sought to identify other factors involved in RMD regulation by surveying effects of siRNAs targeting 54 factors involved in chromatin and the DNA damage response, mismatch repair, and DNA annealing and/or end processing. We measured the effects of siRNAs (pool of 4 per gene) against 54 targets on the frequency of the four RMD events described above, which were compared parallel treatments with a non-targeting siRNA (siCTRL). Each siRNA was tested on all four RMD assays in duplicate, and repeated if the initial fold-effect for any of the assays was ≥ 1.5- fold. We then ranked the results based on the normalized fold-effect at 9.1 kbp for both 1%RMD-GFP and RMD-GFP (Figure 1C). From this analysis, we found siRNAs targeting 21 factors caused a ≥ 1.5-fold effect on at least one of the four RMD assays (Figure 1C, highlighted in red). We then examined these 21 factors using the NHEJ assay, and performed a one-way ANOVA with a Tukey’s post-test to compare the fold-effects between all five assays: the four RMD events and NHEJ. We found that siRNAs targeting 18 of the 21 factors caused a significant difference in at least one RMD event relative to NHEJ, and for all 21 siRNAs we confirmed depletion of the target RNA via qRT-PCR (Figure 2, S2A, S2B).

**Figure 2:**
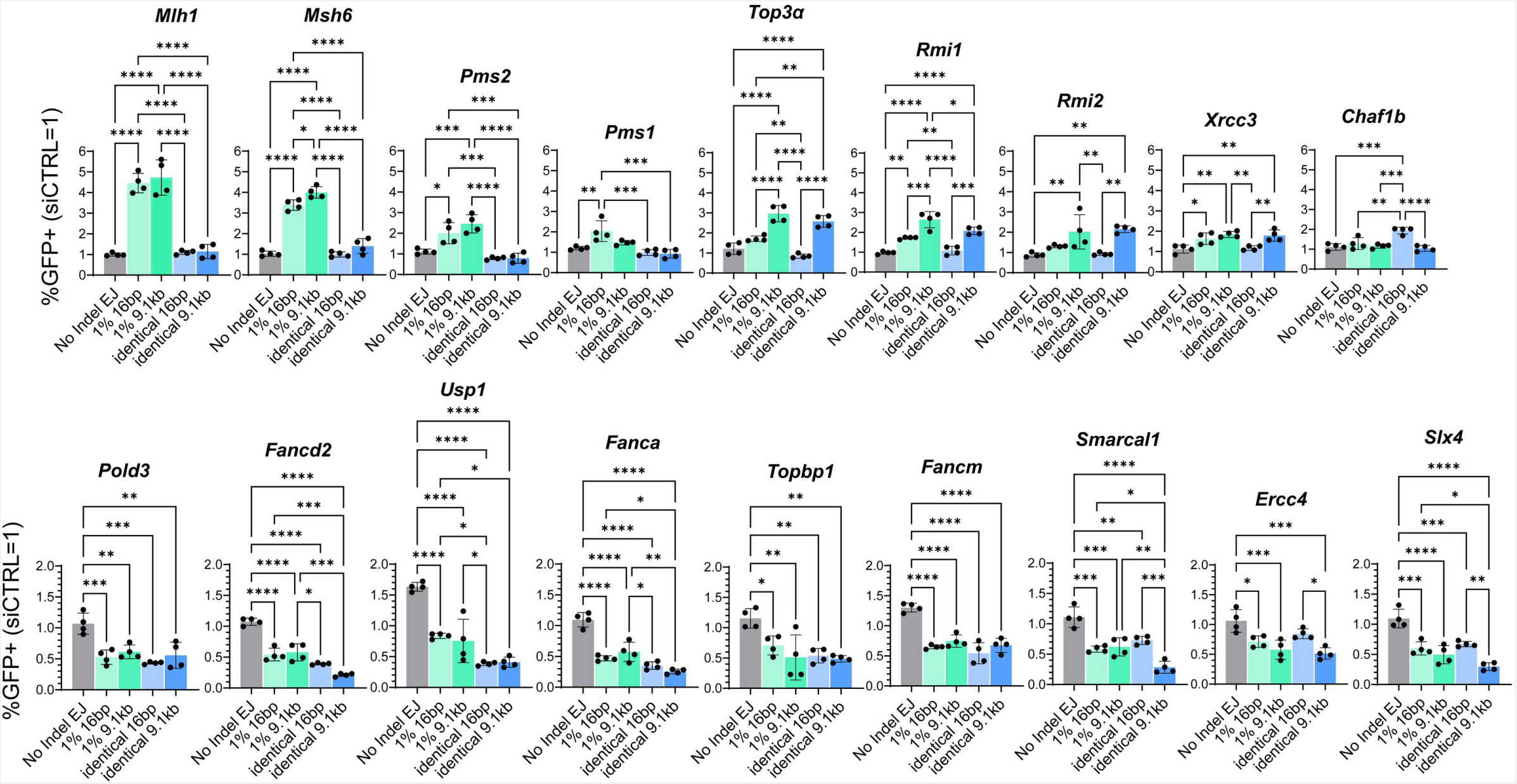
SiRNAs targeting 18 different DNA damage response factors caused a significant difference in at least one RMD event relative to NHEJ. Shown are effects of siRNAs against 18 genes on four RMD events, and NHEJ (No Indel EJ). Frequencies are normalized to transfection efficiency and parallel siCTRL (=1). n=4. *p ≤ 0.05, **p ≤ 0.005, ***p ≤ 0.0005, ****p < 0.0001, one-way ANOVA using Tukey’s multiple comparisons test. Data are represented as mean values ± SD.

The 18 factors fell into different categories based on the relative effects on the distinct RMDs (Figure 2). Nine of the factors, several of which are in the Fanconi Anemia pathway, (POLD3, FANCD2, USP1, FANCA, TOPBP1, FANCM, SMARCAL1, ERCC4, and SLX4) had similar effects as CtIP. Namely, depletion of these factors caused a significant decrease in all four RMD events. Thus, these factors likely mediate steps that are common among all RMDs. In contrast, siRNAs targeting the remaining nine factors (MLH1, MSH6, PMS2, PMS1, TOP3α, RMI1, RMI2, XRCC3, and CHAF1B) caused an increase in at least one RMD event, indicating these factors suppress RMDs. The siRNAs targeting MLH1, MSH6, and PMS2 each caused a significant increase in RMDs with repeat divergence irrespective of DSB/repeat distance, and had no effect on RMDs with identical repeats. Similarly, siRNAs targeting PMS1 caused an increase in RMDs with repeat divergence (1%RMD-GFP) but only for the 16 bp DSB/repeat distance. Thus, these factors (MLH1, MSH6, PMS2, and PMS1) appear to suppress RMDs with repeat divergence. The pattern is more complex with siRNAs targeting TOP3α, RMI1, and RMI2, which are components of the BTR complex (BLM-TOP3α-RMI1/2). Specifically, these factors caused the greatest fold-increases for the 9.1 kbp DSB/repeat distance irrespective of repeat divergence, followed by a more modest increase for 16 bp with 1%RMD-GFP, and no statistical difference for 16 bp with RMD-GFP (Figure 2). Finally, the siRNA targeting CHAF1B caused a specific increase with RMD-GFP at the 16 bp DSB/repeat distance, and conversely targeting XRCC3 caused a modest increase in each of the RMD events except with RMD-GFP at the 16 bp DSB/repeat distance. Altogether, these findings indicate that factors from several pathways, including the Fanconi Anemia pathway, mismatch repair, and the BTR complex influence RMD formation in ways that can be affected by repeat divergence and/or DSB/repeat distance.

### MLH1 suppresses RMDs with divergent repeats

Based on the above survey, we chose to focus on MLH1, both because of its marked effect on the RMDs with divergent repeats, and because its influence on regulation of RMDs, and indeed homologous recombination in mitotic cells, remains poorly understood. For example, while *MLH1* in yeast is critical for mismatch repair, it appears dispensable for suppression of mitotic homologous recombination events (Sugawara et al. 2004; Chakraborty et al. 2016; Pannafino and Alani 2021). We first generated an *Mlh1^-/-^* mESC line by targeting sgRNAs/Cas9 to exon 11 of *Mlh1* that we confirmed has loss of MLH1 by immunoblotting (Figure 3B). We also created an MLH1 expression vector that we validated with immunoblotting (Figure 3B). We then integrated the three RMD reporters (RMD-GFP, 1%RMD-GFP, 3%RMD-GFP) in the *Mlh1^-/-^* mESC line, and these RMD assays were tested using six different 3’ DSB/repeat distances: five that were previously described (16 bp, 3.3 kbp, 9.1 kbp, 19 kbp, 28.4 kbp) (Mendez-Dorantes et al. 2018), whereas the sixth (1 kbp) was added for this study to fill a gap between 16 bp and 3.3 kbp. To validate the 1 kbp DSB site, we used TIDE (tracking of indels by decomposition) analysis (Brinkman et al. 2014), which confirmed induction of indels at the predicted 1 kbp DSB site (Supplemental Figure S3). Also with this TIDE analysis, we found that indel frequencies for the 1 kbp DSB site were similar to the 16 bp and 9.1 kbp DSB sites (Supplemental Figure S3). We compared the results of the RMD assays in the *Mlh1^-/-^*cell lines to WT cells (transfected with empty vector, EV), and also to the complemented condition (*Mlh1^-/-^* transfected with the MLH1 complementation vector) (Figure 3C).

**Figure 3:**
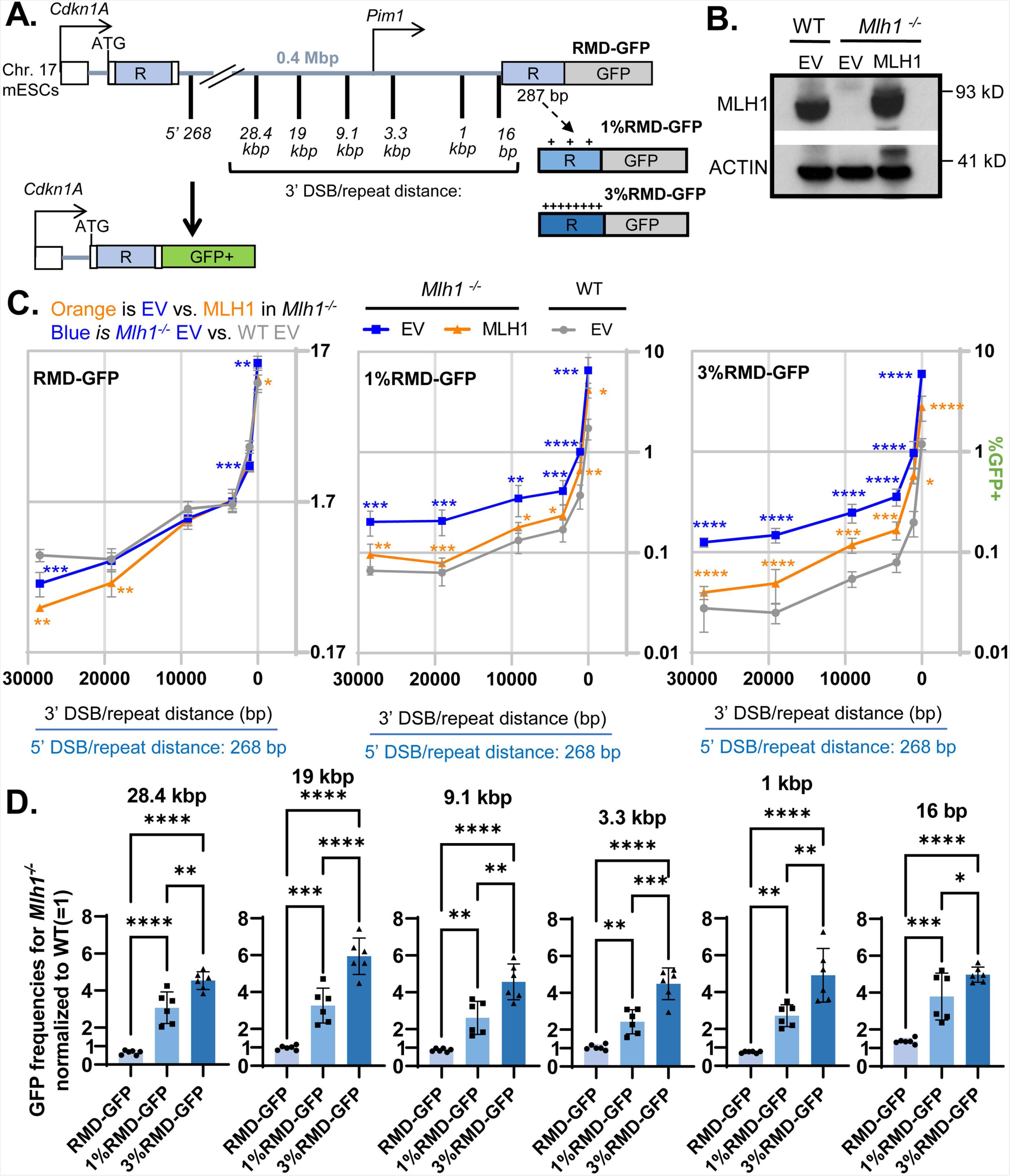
MLH1 suppresses RMDs with divergent repeats. **(A)** Shown is the RMD-GFP reporter as in Figure 1A, but with all DSB sites represented, and with the three repeat sequence versions: identical, 1%RMD-GFP (1% divergence) and 3%RMD-GFP (3% divergence). **(B)** Immunoblotting analysis of MLH1 and ACTIN in WT and *Mlh1^-/-^* mESCs transfected with either EV or MLH1 vectors. **(C)** RMD frequencies for the assays shown in (A), normalized to transfection efficiency, for WT transfected with empty vector (EV), *Mlh1^-/-^* transfected with EV, and *Mlh1^-/-^* transfected with MLH1 expression vector. n=6. *p ≤ 0.05, **p ≤ 0.005, ***p ≤ 0.0005, ****p < 0.0001, unpaired t-test with Holm-Sidak correction. **(D)** Effects of sequence divergence on the relative influence on MLH1 on RMDs. RMD frequencies *Mlh1^-/-^* shown in (C) were normalized to WT (=1), and grouped by location of the DSB upstream of the 3’ repeat to enable comparisons of effects of sequence divergence. n=6. *p ≤ 0.05, **p ≤ 0.005, ***p ≤ 0.0005, ****p < 0.0001, one-way ANOVA with Tukey’s multiple comparisons test. Data are represented as mean values ± SD.

From this analysis, MLH1 showed largely no effect on RMDs between identical repeats, although mild (≤ 1.5-fold) effects were observed at 28.4 kbp, 1 kbp, and 16 bp (*Mlh1^-/-^* vs WT, Figure 3C). However, for RMDs with divergent repeats (1% and 3%), loss of MLH1 caused a significant increase in RMDs at all DSB/repeat distances, both by comparing *Mlh1^-/-^* vs WT, and vs. the complemented cells (*Mlh1^-/-^* cells transfected with the MLH1 expression vector, Figure 3C). We then compared the fold effects of MLH1 loss (*Mlh1^-/-^* vs WT) among the degrees of repeat divergence (identical, 1%, and 3%) for each DSB/repeat distance. Loss of MLH1 caused a significant increase in RMDs at all DSB/repeat distances in 1%RMD-GFP compared to RMD-GFP, and in 3%RMD-GFP compared to 1%RMD-GFP. Thus, the role of MLH1 in suppressing RMDs increased as divergence between the repeats increased (Figure 3D). In contrast, the role of MLH1 was not significantly different between distinct DSB/repeat distances for 1%RMD-GFP and 3%RMD-GFP, although some minor statistical differences based on DSB/repeat distance were observed for RMD-GFP (Supplemental Figure S4). These findings indicate that MLH1 is critical to suppress RMDs if the repeats contain sequence divergence, irrespective of DSB/repeat distance.

### MLH1 and MSH6 function epistatic to each other and MSH2, but independently of EXO1, for suppression of RMDs

Because MLH1 is part of the mismatch repair pathway, we compared its effect to other mismatch repair components, including whether combined mutants show epistatic effects. During mismatch repair, MSH2 and MSH6 form the MUTSα complex and recognize sites of mismatches to then recruit downstream effectors MLH1 and EXO1 for strand nicking and excision (Modrich 2006). To examine the interplay between these factors for RMD regulation, we examined effects of depleting MLH1 and MSH6 in WT, *Mlh1^-/-^*, *Msh2^-/-^*, and *Exo1^-/-^* mESCs. For this analysis, we tested all three RMD reporters (identical repeats, 1 %, and 3% divergent repeats), each at the 16 bp and 9.1 kbp DSB/repeat distances.

We found that depletion of MLH1 in WT and *Exo1^-/-^* mESCs caused a marked increase in RMDs between the divergent repeats at both DSB/repeat distances, but not identical repeats (Figure 4A, B). In contrast, depletion of MLH1 in *Msh2^-/-^* mESCs failed to cause an increase in any of the RMD events tested (Figure 4A, 4B). We found analgous results with MSH6, in that depletion of this factor caused an increase in RMDs between divergent repeats in both WT and *Exo1^-/-^* mESCs, but not in *Mlh1^-/-^*and *Msh2^-/-^* mESCs (Figure 4C, 4D). We confirmed knockdown of MLH1 and MSH6 in each of the genetic backgrounds via immunoblotting (Supplemental Figure S4B, 4C). These results indicate that MLH1 and MSH6 suppress RMDs with divergent repeats independently of EXO1, but function epistatic to each other and to MSH2.

**Figure 4:**
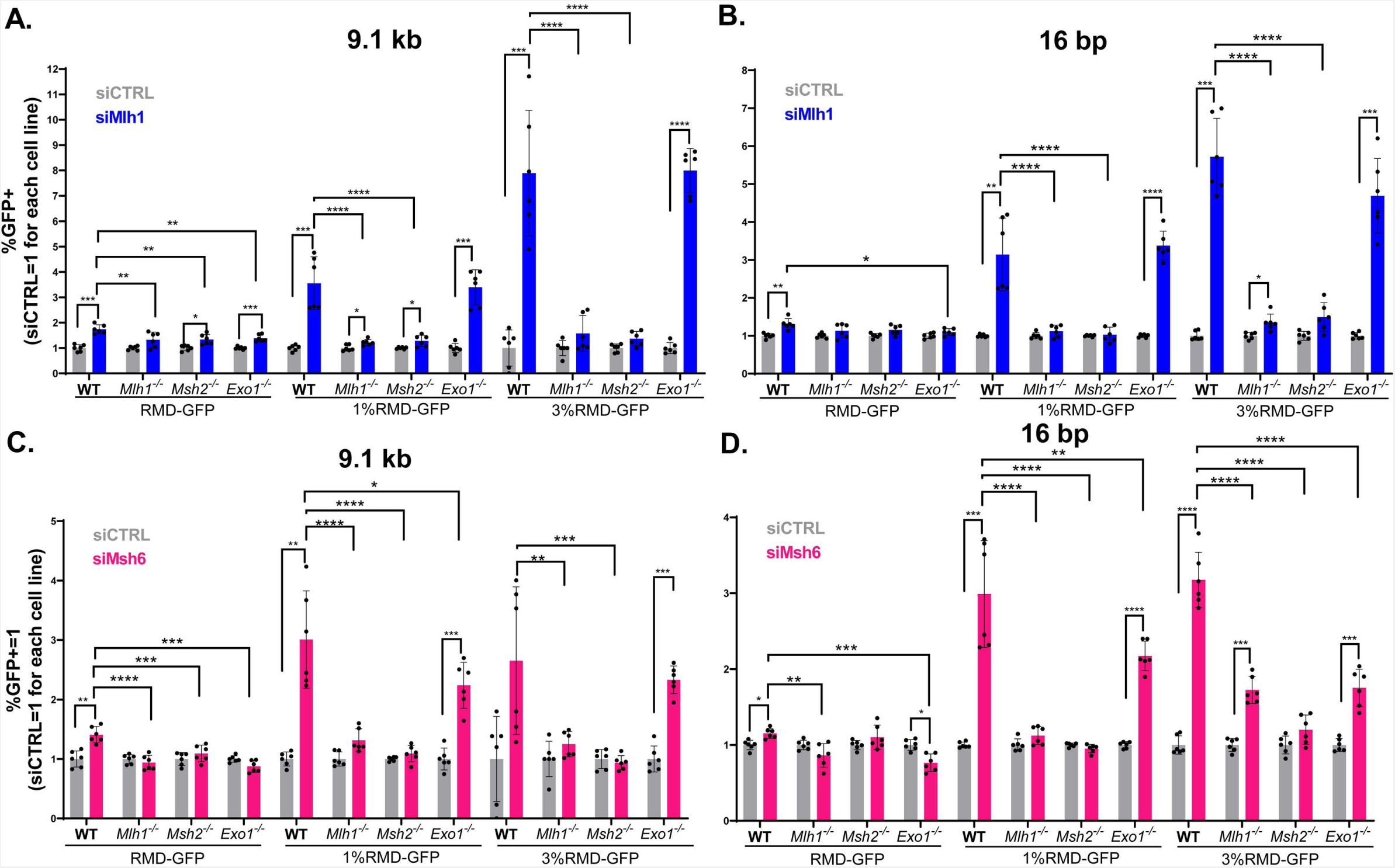
MLH1 and MSH6 are epistatic to each other and MSH2, but independent of EXO1, for suppression of RMDs. **(A)**. Shown is the effect of siRNAs targeting MLH1 (siMlh1) on three RMD events (9.1 kbp DSB/repeat distance, RMD-GFP, 1%GFP-GFP, 3%RMD-GFP) in WT, *Mlh1^-/-^*, *Msh2^-/-^*, and *Exo1^-/-^* mESCs. Frequencies are normalized to transfection efficiency and parallel siCTRL (=1). n=6. *p ≤ 0.05, **p ≤ 0.005, ***p ≤ 0.0005, ****p < 0.0001, unpaired *t*-test for siCTRL vs siMlh1, and unpaired *t*-test using Holm-Sidak correction for effect of siMlh1 in WT vs. the other genetic backgrounds. **(B)** Shown is the analysis as in (A), but using the 16 bp DSB/repeat distance. n=6. Statistics as in (A). **(C)** Shown is the analysis as in (A), but for effects of siRNAs targeting MSH6 (siMsh6). n=6. *p ≤ 0.05, **p ≤ 0.005, ***p ≤ 0.0005, ****p < 0.0001, unpaired *t*-test for siCTRL vs siMsh6, and unpaired *t*-test using Holm-Sidak correction for effect of siMsh6 in WT vs. the other genetic backgrounds **(D)** Shown is the analysis in (C), but using the 16 bp DSB/repeat distance n=6. Statistics as in (C). Data are represented as mean values ± SD.

### The MUTL*α* endonuclease domain is important to suppress RMDs between divergent repeats

We next examined the mechanism by which MLH1 may suppress RMDs. MLH1 interacts with several proteins, including three heterodimer binding partners to form the MUTLα (MLH1:PMS2), MUTLβ (MLH1:PMS1), and MUTLγ (MLH1:MLH3) complexes (Pannafino and Alani 2021). Furthermore, the MUTLα and MUTLγ complexes have endonuclease activity (Harfe et al. 2000; Campbell et al. 2014; Pannafino and Alani 2021). In our siRNA survey described above, we found that siRNAs targeting MLH3 did not affect RMDs, whereas siRNAs targeting PMS2 and PMS1 individually caused a ≥1.5- fold increase in RMDs with divergent repeats (Figure 1C). Thus, we sought to further evaluate the influence of MUTLα (MLH1:PMS2), MUTLβ (MLH1:PMS1), as well as the role of the endonuclease domain of MUTLα (MLH1:PMS2) on RMDs.

To begin with, we tested how siRNAs targeting PMS2 and PMS1 individually, and combined together, affect four distinct RMD events: the two divergent repeat assays (1%RMD-GFP and 3%RMD-GFP), each at two DSB/repeat distances (9.1 kbp and 16 bp). We found that siRNAs targeting PMS1 caused a significant increase in RMDs at both 9.1 kbp and 16 bp in the 1% divergent reporter, and at 16 bp in the 3% reporter, but not at 9.1 kbp in the 3% divergent reporter (Figure 5A). We also found that siRNAs targeting PMS2 caused a significant increase in all four of these RMD events, where the fold-effects were either similar or greater than the effects of siRNAs targeting PMS1 (Figure 5A). Finally, combining siRNAs targeting PMS2 and PMS1 caused the greatest increase in all four of these RMD events that was significantly higher than depleting the two factors alone (Figure 5A). As controls, we also evaluated depletion of PMS2 and PMS1 in the RMD assay with identical repeats and found largely no effect on RMDs (Supplemental Figure S5A). Furthermore, siRNAs targeting PMS2 and PMS1 had no effect on RMD frequencies in the *Mlh1^-/-^* mESCs (all of the identical and divergent repeat assays tested at 16 bp and 9.1 kbp DSB/repeat distance, Supplemental Figure S5B). We confirmed knockdown of PMS2 and PMS1 transcript relative to siCTRL treated cells in both WT and *Mlh1^-/-^* mESCs via qRT-PCR (Supplemental Figure S5C, D). Altogether, these findings indicate that MUTLα and MUTLβ have a role in MLH1-dependent suppression of divergent RMDs.

**Figure 5:**
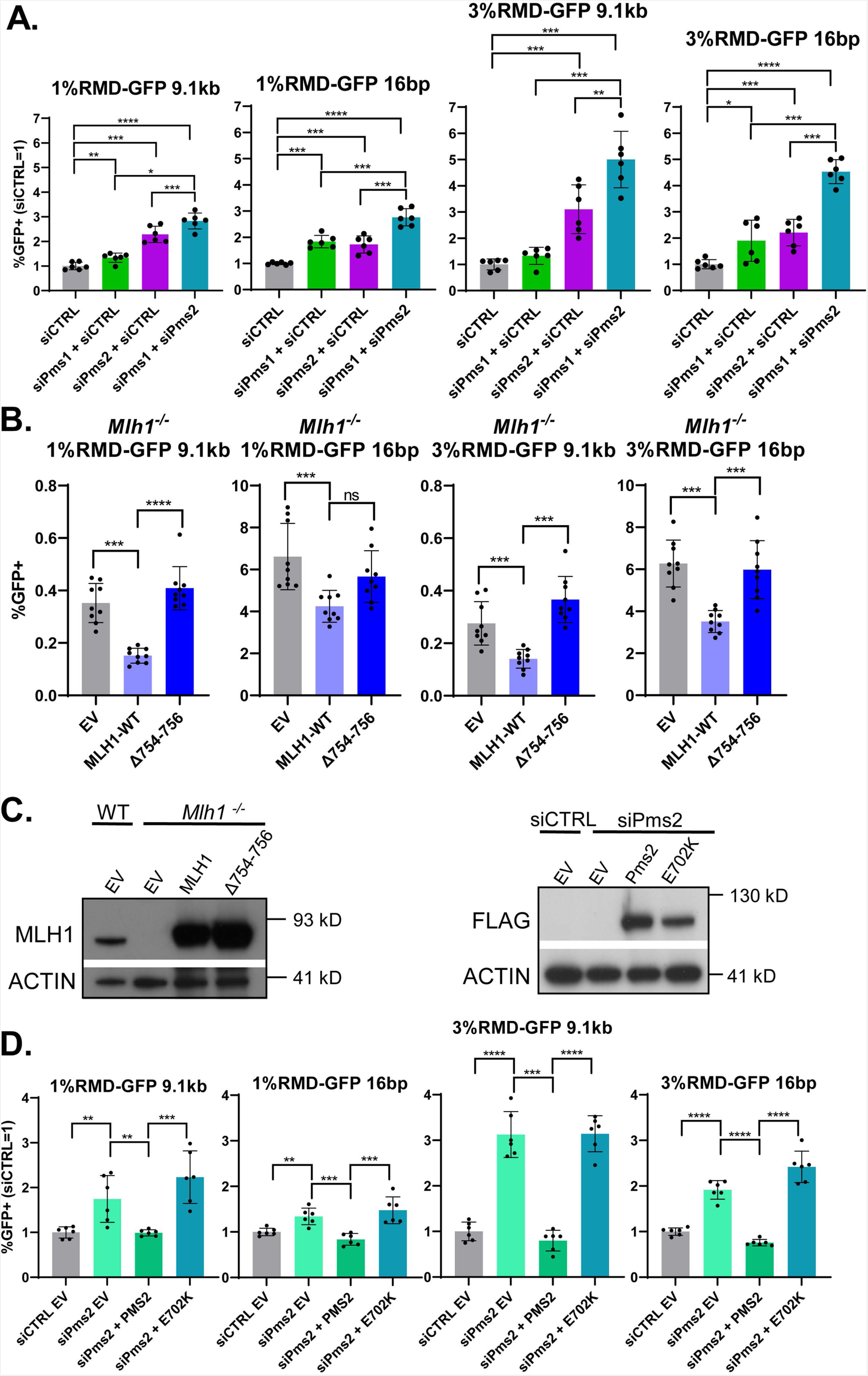
The MUTL*α* endonuclease domain is important to suppress RMDs between divergent repeats. **(A)** Shown are the effects of siRNAs targeting PMS2 (siPms2) and Pms1 (siPms1) individually, and in combination (siPms2 + siPms1), on four RMD events: 1%RMD-GFP, 3%RMD-GFP, each at the 9.1 kbp and 16 bp DSB/repeat distances. siCTRL is added to the siRNA treatments targeting the individual genes to ensure the same total siRNA concentration. Frequencies are normalized to transfection efficiency and parallel siCTRL (=1). n=6. *p ≤ 0.05, **p ≤ 0.005, ***p ≤ 0.0005, ****p < 0.0001, siCTRL vs. each set of siRNA treatments, and also the combination (siPms2 + siPms1) vs. the individual genes, each with unpaired *t*-test using Holm-Sidak correction. **(B)** Shown are RMD frequencies for the four RMDs shown in (A) in *Mlh1^-/-^* mESCs transfected with EV, MLH1-WT, or MLH1-Δ746-756 that deletes the 3 residues at the C-terminus. Frequencies are normalized to transfection efficiency. n=9. ***p ≤ 0.0005, ****p < 0.0001, ns = not significant, unpaired *t*-tests with Holm-Sidak correction. **(C)** Immunoblotting analysis of MLH1 and ACTIN in WT and *Mlh1^-/-^* mESCs transfected with EV, MLH1-WT, or MLH1-Δ746-756 (left). Also shown is immunoblotting analysis of FLAG-PMS2 and ACTIN in WT mESCs transfected with siCTRL EV or siPms2 with EV, PMS2-WT, or PMS2-E702K (right). **(D)** Shown are RMD frequencies in WT mESCs transfected with either siCTRL EV or siPms2 with EV, PMS2-WT, or PMS2-E702K. Frequencies are normalized to transfection efficiency and parallel siCTRL (=1). n=6. **p ≤ 0.005, ***p ≤ 0.0005, ****p < 0.0001, unpaired *t*-test with Holm-Sidak correction. Data are represented as mean values ± SD.

Given that MUTLα has a role in suppressing divergent RMDs, we then considered that its nuclease domain might be important for this function. To test this hypothesis, we examined mutants of MLH1 and PMS2 that have been shown to lack endonuclease activity. We first tested an MLH1 mutant (Δ754-756) with the final three C-terminal amino acids deleted, which have been shown to reside in the metal binding domain that is critical for MUTLα endonuclease activity, but are apparently dispensable for binding to PMS2 (Gueneau et al. 2013; Dai et al. 2021). We then compared RMD frequencies at 16 bp and 9.1 kbp in the two divergent reporters in *Mlh1^-/-^* mESCs expressing either MLH1 WT or Δ754-756. We found that at both 16 bp and 9.1 kbp, Δ754-756 failed to reduce RMDs (Figure 5B). We confirmed both MLH1 WT and Δ754-756 expression via immunoblot (Figure 5C). We next tested a PMS2 endonuclease deficient mutant (E702K), which also disrupts the metal binding domain of MUTLα (van Oers et al. 2010). Specifically, we expressed siRNA resistant forms of PMS2 WT and E702K in cells treated with the siRNAs targeting PMS2. We examined the same four RMD events described above, and found that expression of PMS2 WT, but not E702K, inhibits RMDs between divergent repeats (Figure 5C, D). We also confirmed PMS2 WT and E702K expression via immunoblotting using a 3xFLAG immunotag (Figure 5C). In summary, these findings indicate that the endonuclease domain of MUTLα is important for suppression of RMDs between divergent repeats.

### Mismatch resolution in RMDs with divergent repeats exhibits polarity that is mediated by the MLH1 endonuclease

We next considered that the MLH1 endonuclease might also influence the resolution of the RMD product. Specifically, we considered whether MLH1 might cleave the heteroduplex intermediate in a manner that affects the pattern of mismatch resolution in the RMD. To address this hypothesis, we first tested whether resolution of mismatches in the RMD product follows a specific pattern, or is random. We used the 3%RMD-GFP reporter to determine which base for each of the 8 mismatches was retained in the final RMD product (Figure 6A). We performed this reporter assay using two different 3’ DSB/repeat distances (16 bp and 1 kbp) in WT mESCs, sorted the GFP+ cells by flow cytometry, amplified the rearrangements, and performed deep sequencing analysis. Each of the 8 mismatches were scored as having either the base from the 5’ repeat in the *Cdkn1A* locus (top strand), or from the 3’ repeat fused to GFP (bottom strand) (Figure 6A). We numbered the mismatches 1-8 starting from the *Cdkn1A* side. We performed this analysis with three independent transfections/sorts for each condition to determine the mean/standard deviation for the frequency of retention of the base in the top strand.

**Figure 6:**
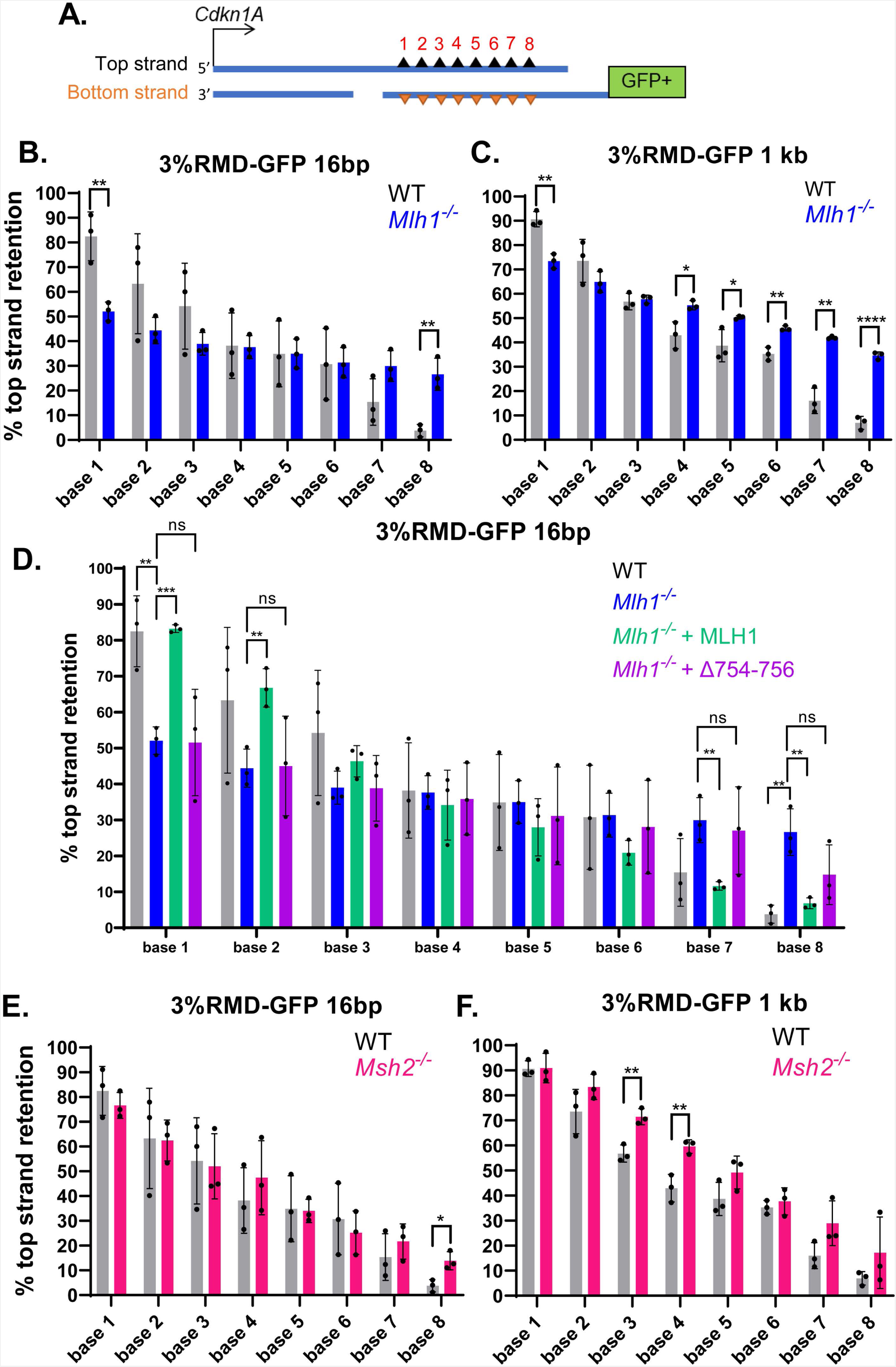
Mismatch resolution in RMDs with divergent repeats exhibits a polarity that is mediated by the MLH1 endonuclease. **(A)** Shown is a diagram of an RMD annealing intermediate with the 8 mismatches of the top and bottom strand shown as triangles, which are numbered 1-8. The DSB ends are shown as DNA nicks without 3’ non-homologous tails only for simplicity. Mismatch resolution was determined by sorting GFP+ cells from the 3%RMD-GFP reporter, amplifying the repeat sequence, and performing deep sequencing analysis to determine the frequency of retention of the top strand base for each mismatch. **(B)** Shown is the frequency of top strand base retention for WT and *Mlh1^-/-^* mESCs using the 16 bp DSB/repeat distance. n=3. **p ≤ 0.005, unpaired *t*-test. **(C)** Shown is the analysis as in (B), except using the 1 kbp DSB/repeat distance. n=3. *p ≤ 0.05, **p ≤ 0.005, ****p < 0.0001, unpaired *t*-test. **(D)** Shown is the frequency of top strand base retention using the 16 bp DSB/repeat distance for WT, *Mlh1^-/-^*, and *Mlh1^-/-^* transfected with expression vectors for MLH1 (WT) and MLH1-Δ754-756. Results for WT and *Mlh1^-/-^*are the same as in (B). n=3. **p ≤ 0.005, ***p ≤ 0.0005, ns = not significant, *Mlh1^-/-^* vs. *Mlh1^-/-^* + MLH1 (WT), and *Mlh1^-/-^* vs. *Mlh1^-/-^* + MLH1-Δ754-756, unpaired *t*-test with Holm-Sidak correction. **(E)** Shown is the frequency of top strand base retention at 16 bp in WT and *Msh2^-/-^* mESCs. WT results are the same as in (A). n=3. *p ≤ 0.05, unpaired *t*-test. **(F)** Shown is the analysis as in (E), but using the 1 kbp DSB/repeat distance. WT results are the same as in (B). n=3. **p ≤ 0.005, unpaired *t*-test. Data are represented as mean values ± SD.

We found that the retention of the top strand base showed a striking polarity in WT cells, for both the 16 bp and 1 kbp 3’ DSB/repeat distances (Figure 6B, C). Specifically, on the *Cdkn1A* side there is preferential retention for the top strand base, whereas on the GFP side there is a preferential loss of the top strand base, and the bases in the middle show no strong bias for either base (Figure 6B, C). This polarity is supported by statistical comparisons (Supplemental Table S1). For example, the first base on the *Cdkn1A* side (base 1) shows significantly greater retention of the top strand, compared to bases 4 through 8. Conversely, the last base from the *Cdkn1A* side (base 8) shows significantly lower retention of the top strand, compared to bases 1 through 3. Based on the SSA model for RMDs, this pattern is consistent with preferential loss of the mismatched bases nearest to the DSB end (Figure 6A).

We then examined the RMD mismatch resolution pattern in the *Mlh1^-/-^* cell line, also with both the 16 bp and 1 kbp DSB/repeat distances. We found that while the mismatch resolution polarity is still detectable, the degree of this polarity is markedly reduced, compared to WT (Figure 6B, C, Supplemental Table S1). For example, for both DSB/repeat distances in *Mlh1^-/-^* cells, base 1 exhibits higher strand retention vs. bases 4-6, which was similar to WT (Supplemental Table S1). However, the frequency of top strand retention for base 1 was substantially lower for *Mlh1^-/-^* vs. WT (Figure 6B, C). Conversely, the frequency of top strand retention for base 8 was substantially higher for *Mlh1^-/-^* vs. WT at both DSB/repeat distances (Figure 6B, C, Supplemental Table S1). These data indicate that MLH1 promotes the polarity for mismatch resolution of RMDs between divergent repeats.

We next posited that the endonuclease domain of MLH1 (i.e., residues 754-756, as described above) is important for the polarity for mismatch resolution. To test this hypothesis, we performed the 3%RMD-GFP assay with the 16 bp DSB/repeat distance in *Mlh1^-/-^*mESCs with expression of MLH1-WT and MLH1-Δ754-756, and then examined the sequence of the RMD products, as described above. From these experiments, we found that expression of MLH1-WT, but not MLH1-Δ754-756, caused an increase in top strand retention for bases 1 and 2, and a converse reduction in top strand retention for bases 7 and 8 (Figure 6D). Thus, MLH1-WT expression, but not MLH1-Δ754-756, restored the polarity for mismatch resolution, indicating that the endonuclease domain of MLH1 is important for this polarity.

As MLH1 and MSH2 function epistatically for RMD suppression (Figure 4A), we also examined mismatch resolution in *Msh2^-/-^* mESCs at both 16 bp and 1 kbp. We found that *Msh2^-/-^* vs. WT cells showed very few statistical differences for the frequency of top strand retention (Figure 6E, F). For the 16 bp DSB/repeat distance, only base 8 showed a statistical difference, with *Msh2^-/-^* mESCs showing an increase in top strand retention. Also, for the 1 kbp DSB/repeat distance, bases 3 and 4 showed statistically higher retention of the top strand in *Msh2^-/-^* mESCs vs. WT, which indicates that for these bases, the polarity was enhanced by loss of MSH2. In summary, WT cells have a polarity for mismatch resolution in RMD products between divergent repeats, which is markedly reduced with loss of MLH1 or its endonuclease domain, but is not obviously affected by MSH2.

### TOP3*α* suppresses RMDs in a manner that is distinct from MLH1

Finally, we sought to contrast MLH1 with another factor that we identified in the siRNA survey as also suppressing RMDs: TOP3α. We performed each of the RMD assays with cells treated with siRNAs targeting TOP3α, which were also co-transfected with either a TOP3α expression vector with silent mutations to be siRNA-resistant, or EV (Figure 7A). Beginning with RMD-GFP, we found that depletion of TOP3α lead to an increase in RMD events at all DSB/repeat distances except 16 bp (Figure 7A). Furthermore, expression of siRNA resistant TOP3α caused a decrease these events at all DSB/repeat distances except 28.4 kbp. In both the divergent reporters (1% and 3%), disruption of TOP3α caused a significant increase in RMDs at all DSB/repeat distances except 28.4 kbp, and these effects were reversed with the TOP3α expression vector, except for the 19 kbp DSB with 3%RMD-GFP (Figure 7A).

**Figure 7:**
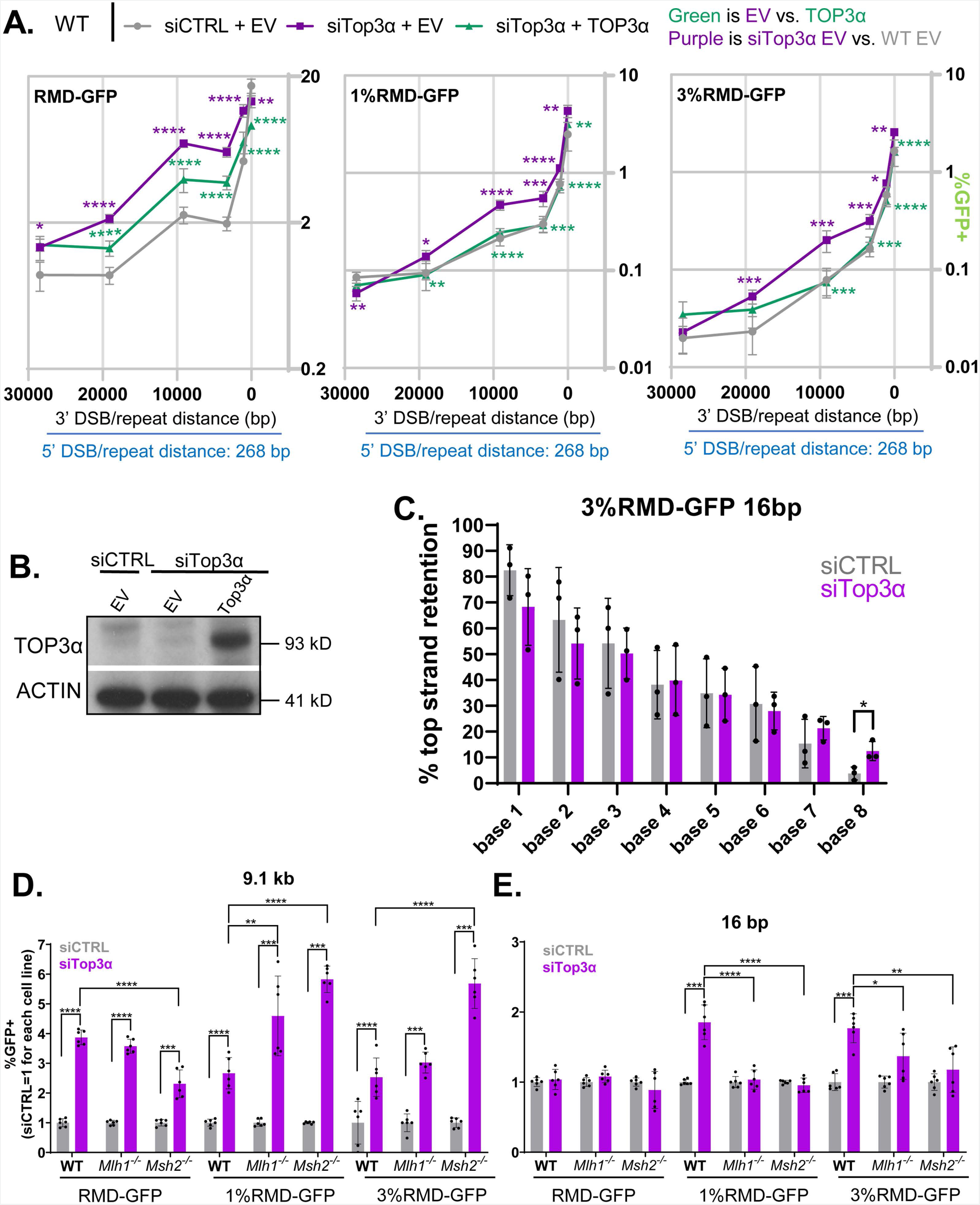
TOP3*α* suppresses RMDs in a manner that is distinct from MLH1. **(A)** Shown are the frequencies of the RMD events depicted in the diagram in Figure 2A (i.e., 6 different DSB/repeat distances with RMD-GFP, 1%RMD-GFP, and 3%RMD-GFP) for WT mESCs transfected with siCTRL and EV, siTop3a and EV, and siTop3α and TOP3α expression vector. n=6. *p ≤ 0.05, **p ≤ 0.005, ***p ≤ 0.0005, ****p < 0.0001, unpaired t-test with Holm-Sidak correction. **(B)** Immunoblotting analysis of TOP3α and ACTIN in WT mESCs transfected with either siCTRL EV, siTop3a EV, or siTop3a with TOP3α expression vector. Endogenous mouse TOP3α was not detected, likely due to the immunogen being human TOP3α. **(C)** Shown is the frequency of top strand base retention performed as in Figure 5B, at the 16 bp DSB/repeat distance for WT (siCTRL) and WT siTop3α. WT (siCTRL) values are the same as in Figure 5B. n=3. *p ≤ 0.05, unpaired *t*-test. **(D)** Shown is the effect of siRNAs targeting TOP3α (siTop3α) on three RMD events (9.1 kbp DSB/repeat distance, RMD-GFP, 1%GFP-GFP, 3%RMD-GFP) in WT, *Mlh1^-/-^*, and *Msh2^-/-^*, mESCs. Frequencies are normalized to transfection efficiency and parallel siCTRL (=1). n=6. **p ≤ 0.005, ***p ≤ 0.0005, ****p < 0.0001, unpaired *t*-test for siCTRL vs. siTop3α, and unpaired *t*-test using Holm-Sidak correction for effect of siTop3α in WT vs. the other genetic backgrounds. **(E)** Shown is the analysis as in (D), but using the 16 bp DSB/repeat distance. n=6. Statistics as in (D), except with **p ≤ 0.005. Data are represented as mean values ± SD.

We next confirmed expression of TOP3α using immunoblot analysis (Figure 7B), examined a catalytically dead mutant of TOP3α (Y362F) (Nicholls et al. 2018), and tested effects of TOP3α on mismatch resolution. We found that while TOP3α WT expression can suppress a set of RMDs, the Y362F mutant had no effect (Supplemental Fig S6A). However, with immunoblot analysis, we found that the TOP3α-Y362F mutant had a much lower molecular weight, which is consistent with other reports of TOP3α mutants that are prone to degradation (Supplemental Fig S6B) (Martin et al. 2018). We tested mismatch retention in cells treated with TOP3α siRNA using the 16 bp DSB/repeat distance, finding that mismatch resolution polarity was not obviously affected, with only a slight increase in top strand retention at base 8 (Figure 7C).

Using the RMD frequency data, we then compared the fold-effects of TOP3α depletion among the various degrees of repeat divergence (identical, 1%, and 3%), and for each DSB/repeat distance. We found that RMDs with identical repeats (RMD-GFP) were effected to at least the same degree as the divergent repeat RMDs by TOP3α depletion, except for the 16 bp 3’ DSB/repeat distance (Supplemental Figure S7A). Namely, with the 16 bp DSB, depletion of TOP3α lead to a significant increase in RMD events for the divergent repeats, but not for the identical repeats. With regards to effect of DSB/repeat distance, we found that TOP3α depletion caused different fold-effects dependent on DSB/repeat distance (Supplemental Figure S7B). The most striking difference is with the RMD-GFP assay, for which TOP3α depletion caused a marked increase in RMDs at both 9.1 and 3.3 kbp, which was statistically higher than 19.1 kbp and 1 kbp, which themselves were statistically higher than 28.4 kbp and 16 bp (Supplemental Figure S7B). The effects of DSB/repeat distance with the divergent repeat RMDs was similar, but more modest (Supplemental Figure S7B).

The above findings indicate that the types of RMDs suppressed by TOP3α are distinct from those of MLH1, which led us to hypothesize that loss of these factors may function independently for RMD suppression. Thus, we examined whether depletion of TOP3α caused further increases in RMDs in *Mlh1^-/-^*mESCs for the 16 bp and 9.1 kbp DSB/repeat distances. We also tested *Msh2^-/-^* for comparison. Depletion of TOP3α caused a marked increase in RMDs with the 9.1 kbp DSB in WT, *Mlh1^-/-^*, and *Msh2^-/-^* mESCs, for both identical and divergent repeats (Figure 7D). Interestingly, with the 16 bp DSB and with divergent repeats, depletion of TOP3α only caused an increase in RMDs in WT mESCs, but failed to do so in *Mlh1^-/-^*, and *Msh2^-/-^* (Figure 7E). We confirmed depletion of the TOP3α RNA in each of the cell lines (Supplemental Figure S7C). These results indicate that TOP3α suppresses RMDs in a manner that is not epistatic to MLH1 and MSH2 when the DSB/repeat distance is long, but is epistatic when the DSB/repeat distance is short.

## DISCUSSION

To characterize factors that regulate RMDs, we began with a survey of several DNA damage response factors in mouse cells, and identified 18 different factors that affect the frequency of RMDs, including several mismatch repair factors and components of the BTR complex that suppress RMDs. We then focused largely on MLH1, which we found suppresses RMDs, but only when the repeats contained sequence divergence. Indeed, the fold-suppression of RMDs via MLH1 increases along with sequence divergence. We also found that MLH1 acts epistatically to the MUTSα complex (MSH2 and MSH6), and two MLH1 binding partners (PMS2 and PMS1) for suppression of such RMDs. Finally, we found that the endonuclease domain of the MUTLα complex (MLH1-PMS2) is important to suppress such RMDs. These results are distinct from findings in *S. cerevisiae*, in that MLH1 is critical for mismatch repair, but apparently not suppression of SSA events with divergent repeats (Sugawara et al. 2004; Goldfarb and Alani 2005; Chakraborty and Alani 2016). Although, our findings are consistent with a recent study that the MLH1 endonuclease domain suppresses prime editing in human cells (i.e., recombination events induced by a DNA nick that use a localized reverse transcribed DNA template for gene editing) (Chen et al. 2021).

Apart from suppression of RMDs, we also found that MLH1 is important for the pattern of mismatch resolution at RMDs. Specifically, in WT cells we found a polarity for mismatch resolution involving preferential removal of the mismatched base proximal to each DSB end (Figure 8A). Conversely, we found preferential retention of the mismatched base distal from each DSB end. These findings appear consistent with reports that Alu-Alu RMDs show polarity in recombination junctions. Namely, the recombination junction for Alu-Alu RMDs are biased towards the 5’ end of Alu elements, which is likely distal to the DSB end (Sen et al. 2006; Morales et al. 2015; Morales et al. 2021). It is unclear whether the mismatch resolution polarity phenomenon described here is related to the recombination junction polarity observed with Alu-Alu RMDs.

**Figure 8:**
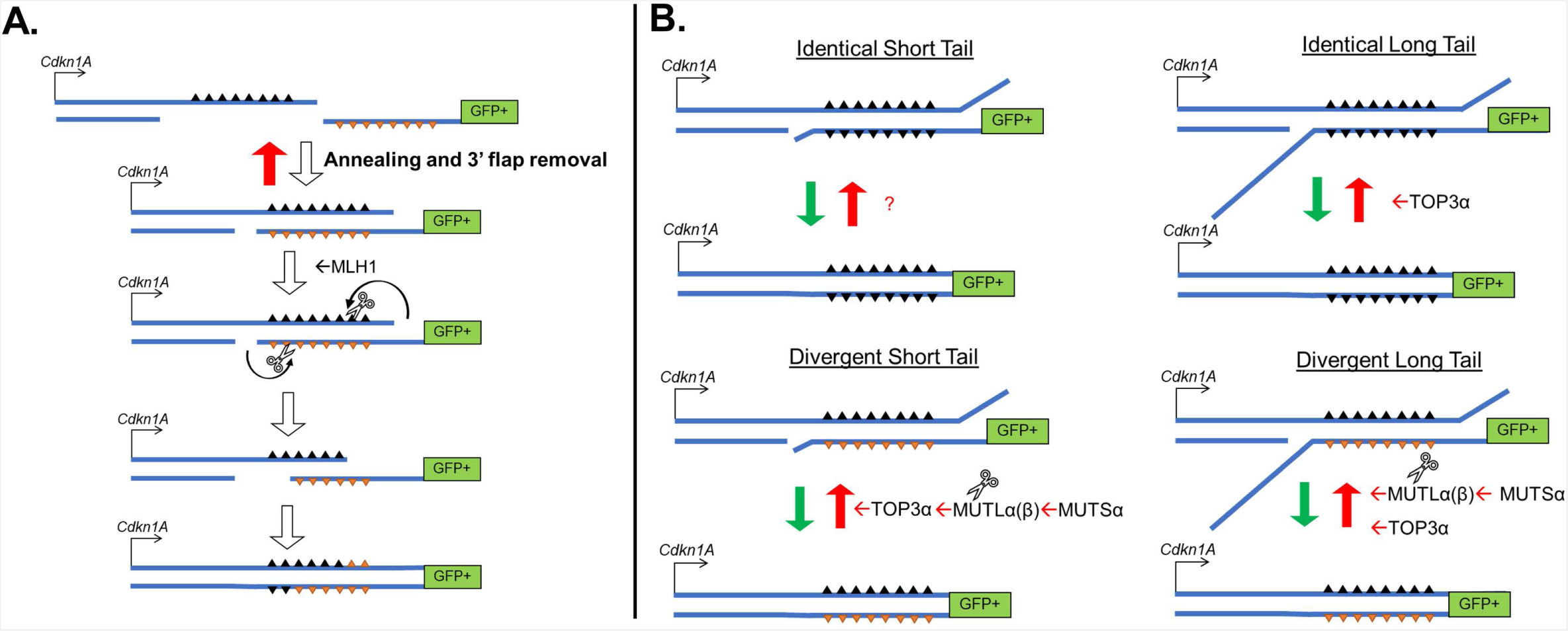
Model. **(A)** Model for the polarity of mismatch resolution in RMDs, which is promoted by the MLH1 endonuclease. **(B)** Model for RMD suppression for short and long DSB/repeat distances (and hence short vs. long 3’ non-homologous tails, respectively), with and without sequence divergence between repeats.

We found that the polarity for mismatch resolution is largely dependent on MLH1 and its endonuclease domain (i.e., the polarity failed to be restored with the MLH1Δ754-756 mutant). We propose a model whereby MLH1 creates an incision upstream of mismatched bases on the strand that is proximal to a DSB end, which initiates degradation and/or replication displacement of the incised strand, and hence loss of the mismatched bases on the strand proximal to the DSB end (Figure 8A). The bias towards creating an incision proximal to the DSB end could be due to the presence of a free DNA end, or DNA nick, on this strand, which is similar to models of mismatch repair at the replication fork. Specifically, components of the replisome (i.e., PCNA and RFC) and MUTSα appear to direct MUTLα to cleave nicked heteroduplex DNA with a strand bias to the nicked DNA strand (Pluciennik et al. 2010; Genschel et al. 2017). These studies with purified proteins support a model of replisome-directed incision of heteroduplex DNA via MUTLα that is biased to the nascent strand due to the presence of a DNA nick at the 3’ end of the nascent strand. Consistent with this model, overexpression of DNA ligase in *S. cerevisiae* causes an increase in mutation rates, and hence reduced mismatch repair, which appears to be caused by premature loss of the DNA nick on the nascent strand (Reyes et al. 2021). While this polarity for mismatch repair with purified proteins is consistent with the polarity we observe with RMDs in mouse cells, the mechanisms may not be precisely the same.

Indeed, there are apparent distinctions between mismatch resolution with purified proteins vs. the RMDs measured in our study, due the findings with MSH2. Namely, while MSH2 is important to direct MUTLα to cleave nicked heteroduplex DNA (Pluciennik et al. 2010; Genschel et al. 2017), we did not find an obvious role for MSH2 in the polarity of mismatch resolution for RMDs. We speculate that MLH1 may be directly recruited to DNA nicks or 3’ non-homologous tails to cleave upstream from mismatched bases that are proximal to the DSB end (Figure 7A). These findings are consistent with the intrinsic nuclease activity of MLH1 in complex with PMS2 (i.e., MUTLα), although certainly this activity is markedly activated with inclusion of other factors (e.g. MUTSα, PCNA, and RFC) (Kadyrov et al. 2006; Gueneau et al. 2013). Another implication of these findings is that suppression of RMDs between divergent repeats vs. mismatch resolution appear to have distinct mechanisms. Namely, as mentioned above, the effects of MLH1 in suppressing RMDs between divergent repeats are epistatic with MUTSα (i.e., MSH2 and MSH6). Altogether, we suggest that MLH1 has multiple roles in regulation of RMDs between divergent repeats: both suppression of these events in a manner that is epistatic with MUTSα, and also an independent role in mismatch resolution to promote the preferential loss of the mismatched base near the chromosomal break end.

Finally, we also found marked distinctions between MLH1 vs. TOP3α in suppression of RMDs. For one, nearly all of the RMD events we examined are suppressed by TOP3α, largely irrespective of sequence divergence or DSB/repeat distance. Interesting exceptions include the short DSB/repeat distance (16 bp) for RMDs with identical repeats, as well as RMDs with the longest DSB/repeat distance (28.4 kbp). Accordingly, TOP3α appears to have a relatively promiscuous anti-RMD activity, which is distinct from the influence of MLH1, which is dependent on sequence divergence. Furthermore, the effects of TOP3α were independent of MLH1 (and MSH2) for the 9.1 kbp DSB/repeat distance. However, interestingly TOP3α appears to function epistatically with MLH1 and MSH2 for suppressing divergent RMDs with a short DSB/repeat distance of 16 bp. A likely consequence of a short DSB/repeat distance is the lack of a long non-homologous 3’ tail in the annealing intermediate during SSA. Thus, we suggest that TOP3α functions independently of mismatch repair to suppress RMDs when there is a long 3’ non-homologous tail (e.g. 9.1 kbp DSB/repeat distance), but is mediated by mismatch repair with a short tail (16 bp DSB/repeat distance, and only with sequence divergence) (Figure 8B).

The role of TOP3α in suppressing RMDs is likely linked to its role in the BTR complex, since depletion of BLM, RMI1, and RMI2 each had similar effects (e.g., each suppress RMDs with identical repeats with the 9.1 kbp DSB/repeat distance, but not 16 bp). The BTR complex has been shown to resolve diverse DNA structures (Bizard and Hickson 2020), which likely accounts for its robust anti-RMD activity. The catalytic activity of TOP3α may also be important for suppressing RMDs, but our experiments with the Y362F mutant were inconclusive because the mutant protein migrates at a lower molecular weight, which is consistent with a report that mutants of TOP3α are prone to degradation (Martin et al. 2018). In summary, whereas MLH1 specifically suppresses RMDs between divergent repeats and also mediates the polarity of mismatch resolution, TOP3α suppresses a diverse set of RMDs, which is epistatic with MLH1 and MSH2 when the repeats have sequence divergence, and the DSB/repeat distance is short.

## METHODS

### Oligonucleotides, plasmids, and cell lines

The siRNAs were pools of 4 per gene in equal concentrations, which were from Dharmacon, with the catalog numbers and sequences in Supplemental Table S2. The non-targeting siRNA (siCTRL) was Dharmacon #D001810-01 5’- UGGUUUACAUGUCGACUAA. Other oligonucleotides are in Supplemental Table S3. The reporter plasmids RMD-GFP, 1%RMD-GFP, 3%RMD-GFP were previously described (Mendez-Dorantes et al. 2018)). All sgRNA/Cas9 plasmids used the px330 plasmid (Addgene 42230, deposited by Dr. Feng Zhang) (Ran et al. 2013). The sgRNA sequences for inducing DSBs in the reporters were previously described (Mendez-Dorantes et al. 2018), apart from the 1 kbp DSB, which is in Supplemental Table S3. The plasmids pCAGGS-NZE-GFP (GFP expression vector), pgk-puro, and pCAGGS-BSKX empty vector (EV) were described previously (Gunn and Stark 2012). The expression vectors for MLH1, TOP3α, and PMS2 were generated with gBLOCK (Integrated DNA Technologies) insertions into pCAGGS-BSKX, with the latter two including silent mutations to mutate all four siRNA target sequences. The mutant forms of TOP3α (Y362F) and PMS2 (E702K) were also generated with gBLOCKs, whereas for the MLH1 mutant (Δ754-756) PCR was used to create a fragment with this deletion.

Several mESC lines with the RMD reporters were previously described: WT (Mendez-Dorantes et al., 2018), *Msh2^-/-^* (Claij and te Riele, 2004; Mendez-Dorantes et al., 2018), and *Exo1^-/-^* (Chen et al., 2017; Mendez-Dorantes et al., 2020). The *Mlh1^-/-^* mESC line was derived using two Cas9-mediated DSBs to introduce a deletion in *Mlh1* using the following sgRNAs, cloned into px330: 5’ CATTGACGTCCACGTTCTGA and 5’ CGAAGTTCACTTTCTGCACG. WT mESCs were transfected with these plasmids and pgk-puro using Lipofectamine 2000 (Thermofisher), transfected cells were enriched using transient puromycin (Sigma Adrich) treatment, followed by plating at low density to isolate and screen individual clones for loss of MLH1. The reporter plasmids were integrated into the *Pim1* locus of mESCs using electroporation of linearized plasmid, selection in hygromycin, and screening by PCR, as described previously (Bennardo et al., 2008).

### DSB Reporter Assays

For the RMD assays including siRNA, mESC cells were seeded on a mixture of 3.75 pmol of siRNA using RNAiMAX (Thermofisher) at a cell density of 0.5 × 10^5^ cells per well of a 24-well plate, with 0.5 ml of antibiotic-free media. The next day, each well was transfected with 200 ng of each sgRNA/Cas9 plasmid plus 3.75 pmol of siRNA using Lipofectamine 2000 (Thermofisher), with 0.5 ml of antibiotic-free media. For the RMD assays with expression vectors for various genes, transfections included 200 ng of these vectors, or the EV control (pCAGGS-BSKX). For the EJ7-GFP assay for NHEJ (No Indel EJ), cells were seeded in the same conditions as the RMD reporters, using the two sgRNAs for this assay, as described (Bhargava et al. 2018). Each experiment included parallel transfections with the GFP expression vector, along with the respective expression vectors and/or siRNAs, to normalize all repair frequencies to transfection efficiency. For all reporter assays, three days after transfection, cells were analyzed by flow cytometry using a CyAn-ADP or ACEA Quanteon, as described (Gunn and Stark, 2012).

### Mismatch Resolution Analysis

For the mismatch resolution analysis with 3%RMD-GFP, the transfection conditions were the same as the frequency analysis described above, and all included siRNA (either siCTRL or siTOP3α), except all amounts were scaled at 2-fold to a 12-well dish, and three days after transfection cells were expanded prior to sorting for GFP+ cells, which were cultured for sorting a second time (BD Aria). Genomic DNA from these samples, purified by phenol/chloroform extraction as described (Gunn and Stark 2012), was used to amplify the repeat sequence using RMDjunct368UPillumina and RMDjunct368DNillumina primers, which include the Illumina adapter sequences. The amplicons were subjected to deep sequencing using the Amplicon-EZ service (AZENTA/GENEWIZ), which includes their SNP/INDEL detection pipeline, which aligned the reads to the bottom strand sequence (Figure 5A) as the reference sequence. All reads that represented ≥0.1% of the total reads for each sample were individually aligned to the reference sequence, and each of the 8 the mismatches were identified as being from either the top or bottom strand (Figure 5A), which was used to calculate the percentage of top strand base retention for each mismatch location. Each cellular condition was examined with three independent transfections and GFP+ sorted samples, and the percentage of retention of the top strand base from the three samples was used to calculate the mean and standard deviation.

### Immunoblotting and quantitative reverse transcription PCR (qRT-PCR)

For immunoblotting analysis, cells were transfected using the same total siRNA and plasmid concentrations as for the reporter assays, but using EV instead of sgRNA/Cas9 plasmids and scaled 2-fold using a 12-well dish. For analysis of siRNA treated cells, following the pre-treatment with siRNA using RNAiMAX (Thermofisher), cells were transfected with pgk-puro plasmid (400 ng), EV (800 ng), and siRNA (7.5 pmol). The next day, cells were re-plated into puromycin and cultured for two days to enrich for transfected cells. Cells were lysed with ELB buffer (250 mM NaCl, 5 mM EDTA, 50 mM Hepes, 0.1% (v/v) Ipegal, and Roche protease inhibitor) with sonication (Qsonica, Q800R). Blots were probed with antibodies for CtIP (Active Motif 61141), BLM (Bethyl Laboratories A300-110A), MLH1 (Abcam, ab92312), MSH6 (Proteintech, 18120-1AP), MSH2 (Bethyl Laboratories, A300-452), EXO1 (Bethyl Laboratories, A302-640A), TOP3α (Proteintech, 14525I-AP), FLAG (Sigma, A8592), and ACTIN (Sigma, A2066). Secondary antibodies (Abcam, ab205719, ab205718). ECL reagent (Amersham Biosciences) was used to develop immunoblotting signals. For quantitative RT-PCR (qRT-PCR) analysis to examine mRNA levels, total RNA was isolated (Qiagen RNAeasy), and reverse transcribed with MMLV-RT (Promega). The RT reactions were amplified with primers for target RNA and ACTIN (Supplemental Table S2) using iTaq Universal SYBR Green (Biorad, 1725120), and quantified on (Biorad CRX Connect Real-Time PCR Detection System, 1855201). Relative levels of each mRNA were determined using the cycle threshold (Ct) value for target mRNA for individual PCR reactions subtracted by Ct value for the ACTIN control (ΔCt value), which was then subtracted from the corresponding ΔCt from siCTRL treated cells (ΔΔCt), which was then used to calculate the 2^-ΔΔCt^ value.

### Tracking of Indels by DEcomposition (TIDE) analysis

WT mESCs were transfected using the same total plasmid concentrations as for the reporter assays, but using sgRNA/Cas9 plasmids and pgk-puro plasmid, and scaled 2-fold using a 12-well dish. The next day, cells were re-plated into puromycin and cultured for two days to enrich for transfected cells. Subsequently, genomic DNA samples were amplified using primers flanking the predicted DSB location, the PCR products were gel purified and analyzed by Sanger sequencing (City of Hope Integrative Genomics Core, Applied Biosystems 3730 DNA Analyzer), which was used for TIDE analysis (Brinkman et al. 2014) to determine the frequency of indels (% INDEL).

## ACKNOWLEDGEMENTS

We thank Carlos Mendez-Dorantes for helpful suggestions. This study was funded in part by the National Cancer Institute of the National Institute of Health: RO1A256989, RO1CA197506, R01CA240392 (J.M.S.); P30CA33572 (City of Hope Core Facilities).

## AUTHOR CONTRIBUTIONS

All authors developed reagents or performed experiments; H.T. and J.M.S. designed experiments and wrote the paper, with input from all authors.

