## Supplementary Materials for "Mismatch resolution for repeat-mediated deletions show a polarity that is mediated by MLH1"

**Supplementary Table S1:** Shown is one-way ANOVA results comparing top strand retention for each mismatch to one another for each strand polarity sample. Purple numbers indicate p-value ≤0.05, orange number indicates p-values ≥0.05.

| Comparison | WT 16bp | WT 1kb | *Mlh1*$^{-/-}$ 16bp | *Mlh1*$^{-/-}$ 1 kb | *Msh2*$^{-/-}$ 16bp | *Msh2*$^{-/-}$ 1 kb | siTop3α 16bp |
|---|---|---|---|---|---|---|---|
| base 1 vs. base 2 | 0.6677 | 0.0165 | 0.6781 | 0.0038 | 0.5654 | 0.9047 | 0.7125 |
| base 1 vs. base 3 | 0.2416 | <0.0001 | 0.1308 | <0.0001 | 0.0658 | 0.0762 | 0.4514 |
| base 1 vs. base 4 | 0.0178 | <0.0001 | 0.0769 | <0.0001 | 0.0202 | 0.0017 | 0.0643 |
| base 1 vs. base 5 | 0.0099 | <0.0001 | 0.025 | <0.0001 | 0.0006 | <0.0001 | 0.019 |
| base 1 vs. base 6 | 0.0048 | <0.0001 | 0.0052 | <0.0001 | <0.0001 | <0.0001 | 0.0045 |
| base 1 vs. base 7 | 0.0003 | <0.0001 | 0.0028 | <0.0001 | <0.0001 | <0.0001 | 0.001 |
| base 1 vs. base 8 | <0.0001 | <0.0001 | 0.0007 | <0.0001 | <0.0001 | <0.0001 | 0.0002 |
| base 2 vs. base 3 | 0.9892 | 0.0196 | 0.9164 | 0.0192 | 0.8413 | 0.5192 | 0.9998 |
| base 2 vs. base 4 | 0.3655 | <0.0001 | 0.7904 | 0.0013 | 0.4924 | 0.0191 | 0.7047 |
| base 2 vs. base 5 | 0.2357 | <0.0001 | 0.4483 | <0.0001 | 0.0241 | 0.0007 | 0.3446 |
| base 2 vs. base 6 | 0.1279 | <0.0001 | 0.1323 | <0.0001 | 0.0024 | <0.0001 | 0.1053 |
| base 2 vs. base 7 | 0.0094 | <0.0001 | 0.076 | <0.0001 | 0.001 | <0.0001 | 0.0247 |
| base 2 vs. base 8 | 0.0012 | <0.0001 | 0.0185 | <0.0001 | 0.0001 | <0.0001 | 0.0033 |
| base 3 vs. base 4 | 0.8224 | 0.0717 | >0.9999 | 0.8406 | 0.9979 | 0.5223 | 0.9136 |
| base 3 vs. base 5 | 0.6585 | 0.0104 | 0.9821 | 0.0125 | 0.2875 | 0.0318 | 0.5913 |
| base 3 vs. base 6 | 0.442 | 0.0022 | 0.682 | 0.0001 | 0.0366 | 0.0008 | 0.2232 |
| base 3 vs. base 7 | 0.0459 | <0.0001 | 0.4978 | <0.0001 | 0.0145 | <0.0001 | 0.0577 |
| base 3 vs. base 8 | 0.006 | <0.0001 | 0.1716 | <0.0001 | 0.0019 | <0.0001 | 0.008 |
| base 4 vs. base 5 | >0.9999 | 0.9662 | 0.9984 | 0.1702 | 0.6202 | 0.6753 | 0.9975 |
| base 4 vs. base 6 | 0.9967 | 0.6249 | 0.8403 | 0.0015 | 0.115 | 0.0358 | 0.8521 |
| base 4 vs. base 7 | 0.4766 | 0.0002 | 0.6743 | <0.0001 | 0.0482 | 0.0021 | 0.4194 |
| base 4 vs. base 8 | 0.0951 | <0.0001 | 0.2751 | <0.0001 | 0.0064 | <0.0001 | 0.0821 |
| base 5 vs. base 6 | >0.9999 | 0.9908 | 0.9898 | 0.2549 | 0.9238 | 0.5583 | 0.994 |
| base 5 vs. base 7 | 0.6535 | 0.0013 | 0.9415 | 0.0048 | 0.7061 | 0.0582 | 0.7855 |
| base 5 vs. base 8 | 0.1603 | <0.0001 | 0.5866 | <0.0001 | 0.187 | 0.0013 | 0.2424 |
| base 6 vs. base 7 | 0.8509 | 0.0061 | >0.9999 | 0.4261 | 0.9996 | 0.8158 | 0.9919 |
| base 6 vs. base 8 | 0.288 | 0.0001 | 0.9568 | 0.0002 | 0.7834 | 0.0544 | 0.6229 |
| base 7 vs. base 8 | 0.9583 | 0.4345 | 0.9938 | 0.0149 | 0.959 | 0.5371 | 0.9626 |

**Supplementary Table S2:** siRNA list

| Pool Catalog Number | Duplex Catalog Number | Gene Symbol | GENE ID | Gene Accession | Sequence |
|---|---|---|---|---|---|
| M-047552-01 | D-047552-01 | Top3a | 21975 | NM_009410 | GAACAUCGGCUUUGAGAUU |
| M-047552-01 | D-047552-02 | Top3a | 21975 | NM_009410 | CCACAAAGAUGGGACCGUA |
| M-047552-01 | D-047552-03 | Top3a | 21975 | NM_009410 | GACAUUUGCUGGCUCAUGA |
| M-047552-01 | D-047552-04 | Top3a | 21975 | NM_009410 | GAGAUGAUCGGCGACUGUA |
| M-058750-01 | D-058750-01 | Ep400 | 75560 | NM_173066 | AAACCGACCUUUCUAUAUU |
| M-058750-01 | D-058750-02 | Ep400 | 75560 | NM_173066 | CAUGAGGAGUUGUAACAUA |
| M-058750-01 | D-058750-03 | Ep400 | 75560 | NM_173066 | GGACGGUUCUCAGGAUAAA |
| M-058750-01 | D-058750-04 | Ep400 | 75560 | NM_173066 | CUAAGAAGCUUGUCAGAAC |
| M-064499-01 | D-064499-01 | Chaf1b | 110749 | NM_028083 | CACCAAAGCUGUCAAUGUU |
| M-064499-01 | D-064499-02 | Chaf1b | 110749 | NM_028083 | GCAAGGAGCCGGAGCAGAU |
| M-064499-01 | D-064499-03 | Chaf1b | 110749 | NM_028083 | UGACAAACACCACUUAUGU |
| M-064499-01 | D-064499-04 | Chaf1b | 110749 | NM_028083 | GGGCAACCGAUGGGAAUUU |
| M-063756-01 | D-063756-21 | Zswim7 | 69747 | XM_001473892 | GCUAGGGAGUUCUGGCAAA |
| M-063756-01 | D-063756-22 | Zswim7 | 69747 | XM_001473892 | CCUCUAUGGUCUUAAUAUU |
| M-063756-01 | D-063756-23 | Zswim7 | 69747 | XM_001473892 | GUAAGCAUCUUCUGGCAAU |
| M-063756-01 | D-063756-24 | Zswim7 | 69747 | XM_001473892 | CCUUCUCAGUGUUACGGAA |
| M-058186-01 | D-058186-01 | Fancm | 104806 | NM_178912 | GAUGAGGACAUACGAUAUA |
| M-058186-01 | D-058186-02 | Fancm | 104806 | NM_178912 | GGAAACAGGGCUUGCAUUC |
| M-058186-01 | D-058186-03 | Fancm | 104806 | NM_178912 | AAGAAAGAAUGCUGGUAUU |
| M-058186-01 | D-058186-04 | Fancm | 104806 | NM_178912 | GGUUACAAGUAGAGAUUUG |
| M-045170-01 | D-045170-01 | Ercc1 | 13870 | NM_007948 | GUGAGGUGAUUCCCGAUUA |
| M-045170-01 | D-045170-02 | Ercc1 | 13870 | NM_007948 | GAACUUCGCCCUUCGUGUG |
| M-045170-01 | D-045170-03 | Ercc1 | 13870 | NM_007948 | GGCGGUACCUGGAGACCUA |
| M-045170-01 | D-045170-04 | Ercc1 | 13870 | NM_007948 | UGUUGAAGUUUGUGCGCAA |
| M-049323-00 | D-049323-01 | Mlh1 | 17350 | NM_026810 | GCGACAAGGUCUACGCUUA |
| M-049323-00 | D-049323-02 | Mlh1 | 17350 | NM_026810 | GAGAUGCUCCGUAACCAUU |
| M-049323-00 | D-049323-03 | Mlh1 | 17350 | NM_026810 | CGGAGAUGCUUGCAGACUA |
| M-049323-00 | D-049323-04 | Mlh1 | 17350 | NM_026810 | CAAGUGAAGAGUACGGAAA |
| M-044082-00 | D-044082-01 | Msh6 | 17688 | NM_010830 | CUAGAAGGAUUCAAAGUAA |
| M-044082-00 | D-044082-02 | Msh6 | 17688 | NM_010830 | GCAUGAGGCUUAUUUAUCU |
| M-044082-00 | D-044082-03 | Msh6 | 17688 | NM_010830 | CGGAGGAGGUUAUUCAGAA |
| M-044082-00 | D-044082-04 | Msh6 | 17688 | NM_010830 | GCUAUAAUGUACGAAGAAA |
| M-048522-01 | D-048522-01 | Pms2 | 18861 | NM_008886 | GGACUCGACUGGUGUUUGA |
| M-048522-01 | D-048522-02 | Pms2 | 18861 | NM_008886 | GAAGUGGCCAGUAGCUUUA |
| M-048522-01 | D-048522-03 | Pms2 | 18861 | NM_008886 | CGUGUAAGCUGCACUAAUC |
| M-048522-01 | D-048522-17 | Pms2 | 18861 | NM_008886 | CCACAAAUGCCAAGCGCUU |
| M-049174-01 | D-049174-01 | Xrcc3 | 74335 | NM_028875 | GAGAAGAUCAGGUUCAGCA |
| M-049174-01 | D-049174-02 | Xrcc3 | 74335 | NM_028875 | CCGAAUUACUGCUGCGGUU |
| M-049174-01 | D-049174-03 | Xrcc3 | 74335 | NM_028875 | AAGACAAUAUGGAGGCCUA |
| M-049174-01 | D-049174-04 | Xrcc3 | 74335 | NM_028875 | UGUUGGAGUGUGUGAGUAA |
| M-054715-00 | D-054715-01 | Pms1 | 227099 | NM_153556 | UAACAGAGAGUCUUCUUAA |
| M-054715-00 | D-054715-02 | Pms1 | 227099 | NM_153556 | GAAUAUCGAACCUGUGAAA |
| M-054715-00 | D-054715-03 | Pms1 | 227099 | NM_153556 | GUACAUAAUAAGGCAGUUA |
| M-054715-00 | D-054715-04 | Pms1 | 227099 | NM_153556 | AAAGUUAAUUCGACGCUAU |
| M-041643-00 | D-041643-01 | Smarcal1 | 54380 | NM_018817 | GGAGUUAGCUUUGCCAUAU |
| M-041643-00 | D-041643-02 | Smarcal1 | 54380 | NM_018817 | GGAAGAACGUUCAACACAU |
| M-041643-00 | D-041643-03 | Smarcal1 | 54380 | NM_018817 | GGGGAUCGCUUCCGGGUAA |
| M-041643-00 | D-041643-04 | Smarcal1 | 54380 | NM_018817 | CAUGUGUCGUCGAGUAUAU |
| M-056921-01 | D-056921-01 | Fancd2 | 211651 | NM_001033244 | GCUAAAGACCGUUCGCUUA |

| | | | | | |
|---|---|---|---|---|---|
| M-056921-01 | D-056921-02 | Fancd2 | 211651 | NM_001033244 | GCAUGUGUGUUUAAAGUAU |
| M-056921-01 | D-056921-03 | Fancd2 | 211651 | NM_001033244 | UCUAUGGACUGGAAGAGUA |
| M-056921-01 | D-056921-04 | Fancd2 | 211651 | NM_001033244 | ACACAAGACUCACCAAGCA |
| M-045003-00 | D-045003-01 | Ercc4 | 50505 | NM_015769 | GGAGCGUGCUUCCGCCAAA |
| M-045003-00 | D-045003-02 | Ercc4 | 50505 | NM_015769 | GCCUGAAGUUGUAGAGAUU |
| M-045003-00 | D-045003-03 | Ercc4 | 50505 | NM_015769 | UCACAACCCGUCACUUGAA |
| M-045003-00 | D-045003-04 | Ercc4 | 50505 | NM_015769 | GCCGAAUACUCGUGGUUGA |
| M-058407-01 | D-058407-01 | Fanca | 14087 | NM_016925 | UGACAGACCCGACCCAAUA |
| M-058407-01 | D-058407-02 | Fanca | 14087 | NM_016925 | GGAGUCAGUUGGCCGUUUG |
| M-058407-01 | D-058407-03 | Fanca | 14087 | NM_016925 | GACUAAGUGUCCCGUGAUU |
| M-058407-01 | D-058407-04 | Fanca | 14087 | NM_016925 | GUCGGUGGAUGAGAUGUUA |
| M-059440-01 | D-059440-01 | Topbp1 | 235559 | NM_176979 | GGAUGAAUCUAUAUACAAG |
| M-059440-01 | D-059440-02 | Topbp1 | 235559 | NM_176979 | CCACGAUGGUUGAAGCUAA |
| M-059440-01 | D-059440-03 | Topbp1 | 235559 | NM_176979 | GCAGACAAUUACCGUGUAU |
| M-059440-01 | D-059440-04 | Topbp1 | 235559 | NM_176979 | GGAGUUUGCCGGCAGUUAA |
| M-057081-01 | D-057081-01 | Slx4 | 52864 | NM_177472 | CAUCUGAACUGUCACAAAU |
| M-057081-01 | D-057081-02 | Slx4 | 52864 | NM_177472 | GAGCUGCGUUUGAGUAUUC |
| M-057081-01 | D-057081-03 | Slx4 | 52864 | NM_177472 | GCGAGAUGCCCACUUCUUA |
| M-057081-01 | D-057081-04 | Slx4 | 52864 | NM_177472 | GCCCUAAGGUUUGGUGUGA |
| M-054618-01 | D-054618-01 | Pif1 | 208084 | NM_172453 | GAGCCUAGGUCGAAACGAA |
| M-054618-01 | D-054618-02 | Pif1 | 208084 | NM_172453 | CCAAUGGAAUCCCAGACUA |
| M-054618-01 | D-054618-03 | Pif1 | 208084 | NM_172453 | GUACGGUUCCUGUGUGGUA |
| M-054618-01 | D-054618-04 | Pif1 | 208084 | NM_172453 | GCCGUGUCCUUCAGUUAAA |
| M-055235-01 | D-055235-01 | Mus81 | 71711 | NM_027877 | GGACAUUGGCGAAACCAGA |
| M-055235-01 | D-055235-02 | Mus81 | 71711 | NM_027877 | GCAUUGCGCUCCCUCCAAC |
| M-055235-01 | D-055235-03 | Mus81 | 71711 | NM_027877 | GUGAAGCGAACCAUGGAUA |
| M-055235-01 | D-055235-04 | Mus81 | 71711 | NM_027877 | GUACGCAAGCUACACGUUG |
| M-053624-01 | D-053624-01 | Eme1 | 268465 | NM_177752 | AAUCAGGAAUGGCAAAUAA |
| M-053624-01 | D-053624-02 | Eme1 | 268465 | NM_177752 | GAGCAAGAACGUCAGAAUU |
| M-053624-01 | D-053624-03 | Eme1 | 268465 | NM_177752 | GCACGGGACUCAUGGUGUC |
| M-053624-01 | D-053624-04 | Eme1 | 268465 | NM_177752 | GGAUCUACCUUCAAAUGAC |
| M-046305-01 | D-046305-01 | Pold3 | 67967 | NM_133692 | AGAACAAGAUCGUGACUUA |
| M-046305-01 | D-046305-02 | Pold3 | 67967 | NM_133692 | UGGCUAAGCUAUACACUAG |
| M-046305-01 | D-046305-03 | Pold3 | 67967 | NM_133692 | CGAGUAGACUUGUCGGAUG |
| M-046305-01 | D-046305-04 | Pold3 | 67967 | NM_133692 | ACGAAAGCGUGUACUGAAA |
| M-059205-01 | D-059205-01 | Phf8 | 320595 | NM_177201 | GAUGAAGAGUGUAUCCAAA |
| M-059205-01 | D-059205-02 | Phf8 | 320595 | NM_177201 | CGGGAUAGCUAUACAGAUU |
| M-059205-01 | D-059205-03 | Phf8 | 320595 | NM_177201 | GGAAUUCUUGAUCUACUUA |
| M-059205-01 | D-059205-04 | Phf8 | 320595 | NM_177201 | GGAAAAGGCGAACCUGUUA |
| M-047716-00 | D-047716-01 | Msh5 | 17687 | NM_013600 | GCAUAGAUCAAGACACUUA |
| M-047716-00 | D-047716-02 | Msh5 | 17687 | NM_013600 | GAAAUCAUACUUCUGCCAA |
| M-047716-00 | D-047716-03 | Msh5 | 17687 | NM_013600 | GGACUACGGCUAUUCGAGA |
| M-047716-00 | D-047716-04 | Msh5 | 17687 | NM_013600 | CCGCAUUACUCUCCCUGUA |
| M-062301-01 | D-062301-01 | Mcm3ap | 54387 | NM_019434 | CAAGACAGCUCACCAGAUA |
| M-062301-01 | D-062301-02 | Mcm3ap | 54387 | NM_019434 | GCAAGAAGCUCGCCGUGAU |
| M-062301-01 | D-062301-03 | Mcm3ap | 54387 | NM_019434 | GCGACAAACUUCCUCAAUU |
| M-062301-01 | D-062301-04 | Mcm3ap | 54387 | NM_019434 | GUAUAAAGGUGGCUCAUGG |
| M-049912-00 | D-049912-01 | Rpa1 | 68275 | NM_026653 | UGAACAAGGUGUAUUACUU |
| M-049912-00 | D-049912-02 | Rpa1 | 68275 | NM_026653 | GGCUAAAGACGCACUAGUA |
| M-049912-00 | D-049912-03 | Rpa1 | 68275 | NM_026653 | GCUGAAGACGUUGGAUUAA |
| M-049912-00 | D-049912-04 | Rpa1 | 68275 | NM_026653 | CGCAUGAUCUUAUCGGCAA |
| M-055247-01 | D-055247-01 | Atrip | 235610 | NM_172774 | ACGCAGAAAUGAAUGAGUU |
| M-055247-01 | D-055247-02 | Atrip | 235610 | NM_172774 | GAGCAACGGACGAACAAAU |
| M-055247-01 | D-055247-03 | Atrip | 235610 | NM_172774 | UCAUAAGGUCCGCCGAUUA |

| | | | | | |
|---|---|---|---|---|---|
| M-055247-01 | D-055247-04 | Atrip | 235610 | NM_172774 | GGUGAGCACGCGGAAUUUA |
| M-056219-01 | D-056219-01 | Smarcad1 | 13990 | NM_007958 | CCACACAUGUUUAGCAGUA |
| M-056219-01 | D-056219-02 | Smarcad1 | 13990 | NM_007958 | GUUUAAAUGUGCUCUGUUA |
| M-056219-01 | D-056219-03 | Smarcad1 | 13990 | NM_007958 | GCAAUGUCAUGAUGCAAUU |
| M-056219-01 | D-056219-04 | Smarcad1 | 13990 | NM_007958 | GCACGUAACCGUUUAUUGC |
| M-040694-01 | D-040694-01 | Arid1a | 93760 | NM_001080819 | GAAAUGACAUGACCUACAA |
| M-040694-01 | D-040694-02 | Arid1a | 93760 | NM_001080819 | CCACCAGGCUACCCAAAUA |
| M-040694-01 | D-040694-03 | Arid1a | 93760 | NM_001080819 | GAAUACCUCUGACAUGAUG |
| M-040694-01 | D-040694-17 | Arid1a | 93760 | NM_001080819 | UUUAUAGUAUGGCGAGUUA |
| M-062392-02 | D-062392-17 | Setd2 | 235626 | NM_001081340 | CGAAAGAGAUCGAAGGCGA |
| M-062392-02 | D-062392-18 | Setd2 | 235626 | NM_001081340 | GGGAAAUCAUCAAGAUCGA |
| M-062392-02 | D-062392-19 | Setd2 | 235626 | NM_001081340 | CUCCAAUUGUGCAGAGUUA |
| M-062392-02 | D-062392-20 | Setd2 | 235626 | NM_001081340 | GAGUGGAGUAUGAGCGGAA |
| M-042994-01 | D-042994-01 | H2afz | 51788 | NM_016750 | ACAAAUCGCUGAUCGGGAA |
| M-042994-01 | D-042994-02 | H2afz | 51788 | NM_016750 | CAUCGACACCUGAAAUCUA |
| M-042994-01 | D-042994-03 | H2afz | 51788 | NM_016750 | GUCAAAAGACUUAAAGGUA |
| M-042994-01 | D-042994-04 | H2afz | 51788 | NM_016750 | UGCUAUACGUGGAGAUGAA |
| M-044077-01 | D-044077-01 | Msh3 | 17686 | NM_010829 | GAACUGCAGUACUUAGAUA |
| M-044077-01 | D-044077-02 | Msh3 | 17686 | NM_010829 | UAUACGAAAUCCACACUUA |
| M-044077-01 | D-044077-03 | Msh3 | 17686 | NM_010829 | AACUGAAACUGCCGCAUUA |
| M-044077-01 | D-044077-04 | Msh3 | 17686 | NM_010829 | GAUCCAAGAGCGUCUAUAC |
| M-046249-01 | D-046249-01 | Msh4 | 55993 | NM_031870 | GCGCAUGCCUGUACUCUUU |
| M-046249-01 | D-046249-02 | Msh4 | 55993 | NM_031870 | GCACCCAAAUCGCUAAAGA |
| M-046249-01 | D-046249-03 | Msh4 | 55993 | NM_031870 | GAACGUUAAUUUCACUACA |
| M-046249-01 | D-046249-04 | Msh4 | 55993 | NM_031870 | AGAGAUAGUGGACGACAUA |
| M-066289-01 | D-066289-09 | Mlh3 | 217716 | NM_175337 | GCUGAGAGCUUAGCCGUUA |
| M-066289-01 | D-066289-10 | Mlh3 | 217716 | NM_175337 | UGACAGGACUUAGCACAUU |
| M-066289-01 | D-066289-11 | Mlh3 | 217716 | NM_175337 | GAGGAACGAUGUAUCUGGA |
| M-066289-01 | D-066289-12 | Mlh3 | 217716 | NM_175337 | CAAAGAAGAUCGCUUAGAA |
| M-062824-01 | D-062824-01 | Mcm9 | 71567 | NM_027830 | GGUACUGGCUGGUGGAAUU |
| M-062824-01 | D-062824-02 | Mcm9 | 71567 | NM_027830 | CGAGCGGGAUUACAUGUGU |
| M-062824-01 | D-062824-03 | Mcm9 | 71567 | NM_027830 | CCACGGUCUAUGAAAGUUA |
| M-062824-01 | D-062824-04 | Mcm9 | 71567 | NM_027830 | AGAUGUACGCUGUGAAGUU |
| M-060852-01 | D-060852-01 | Asf1a | 66403 | NM_025541 | CUACCGAGGUCAAGAAUUU |
| M-060852-01 | D-060852-02 | Asf1a | 66403 | NM_025541 | AGAGUUGGCUACUAUGUAA |
| M-060852-01 | D-060852-03 | Asf1a | 66403 | NM_025541 | UCGAGGACCUGUCUGAAGA |
| M-060852-01 | D-060852-04 | Asf1a | 66403 | NM_025541 | AAUCUACAGUCCCUUCUUU |
| M-046122-01 | D-046122-01 | Asf1b | 66929 | NM_024184 | GGGCUACUAUGUCAACAAU |
| M-046122-01 | D-046122-02 | Asf1b | 66929 | NM_024184 | CAGUUGCACUCCUGUUAAA |
| M-046122-01 | D-046122-03 | Asf1b | 66929 | NM_024184 | GUGAGGAGUUUGAUCAGAU |
| M-046122-01 | D-046122-04 | Asf1b | 66929 | NM_024184 | UCUCUCAGCUACAGCGGAA |
| M-055156-00 | D-055156-01 | Brip1 | 237911 | NM_178309 | GGUUAUGAGUCGUCUUGUA |
| M-055156-00 | D-055156-02 | Brip1 | 237911 | NM_178309 | GUACGGACCAUUGUUCUAA |
| M-055156-00 | D-055156-03 | Brip1 | 237911 | NM_178309 | GACGGCAUUUCAAUAGUAA |
| M-055156-00 | D-055156-04 | Brip1 | 237911 | NM_178309 | CAACUCAAGUCGUGCUCAA |
| M-044778-01 | D-044778-01 | Recql | 19691 | NM_023042 | GCAAAUCCAUGGAGAAUUA |
| M-044778-01 | D-044778-02 | Recql | 19691 | NM_023042 | GUAAAGACGUUUCGUUUGA |
| M-044778-01 | D-044778-03 | Recql | 19691 | NM_023042 | CAAGUGUCGCCGUGUGUUA |
| M-044778-01 | D-044778-04 | Recql | 19691 | NM_023042 | UUAUAAGGCUCUUGGCAUC |
| M-058494-01 | D-058494-01 | Wrn | 22427 | NM_001122822 | GAAUGACUCCUCCUAUAUA |
| M-058494-01 | D-058494-02 | Wrn | 22427 | NM_001122822 | CGAAUCAUCUUGUCCCAUU |
| M-058494-01 | D-058494-03 | Wrn | 22427 | NM_001122822 | GAAAAGUUCCGGUUAUAUA |
| M-058494-01 | D-058494-04 | Wrn | 22427 | NM_001122822 | GAUCGACGGUGUCUCUGAA |
| M-046995-01 | D-046995-01 | Recql4 | 79456 | NM_058214 | CCAAAUCCAUCAACAGUAA |

| | | | | | |
|---|---|---|---|---|---|
| M-046995-01 | D-046995-02 | Recql4 | 79456 | NM_058214 | UGGCAUAGCUGGCGAGUUU |
| M-046995-01 | D-046995-03 | Recql4 | 79456 | NM_058214 | CCUAGACCCUGGUUGGUUA |
| M-046995-01 | D-046995-04 | Recql4 | 79456 | NM_058214 | UAUAUUCGACUCAACAUGA |
| M-045558-00 | D-045558-01 | Recql5 | 170472 | NM_130454 | GGGCCAACCUCUUCUAUGA |
| M-045558-00 | D-045558-02 | Recql5 | 170472 | NM_130454 | GCUAAUGUCCGGUUUGUUG |
| M-045558-00 | D-045558-03 | Recql5 | 170472 | NM_130454 | GGGAACAAACCGUCUGAUA |
| M-045558-00 | D-045558-04 | Recql5 | 170472 | NM_130454 | GAGGGAGGCCGCAGACAUU |
| M-041700-01 | D-041700-01 | Rad51b | 19363 | NM_009014 | GCAAACGGCUUAUGAGUUA |
| M-041700-01 | D-041700-02 | Rad51b | 19363 | NM_009014 | GAUACCAGAUUGUUAACUG |
| M-041700-01 | D-041700-03 | Rad51b | 19363 | NM_009014 | CAAAUUACGACCCAUCUGA |
| M-041700-01 | D-041700-04 | Rad51b | 19363 | NM_009014 | CAUAAUGAUGAGUGUCUUA |
| M-044655-01 | D-044655-01 | Rad51c | 114714 | NM_053269 | ACAGAGAGUUCUUUGAAUU |
| M-044655-01 | D-044655-02 | Rad51c | 114714 | NM_053269 | GAUAAUAGACGGAAUUGCU |
| M-044655-01 | D-044655-03 | Rad51c | 114714 | NM_053269 | CGUACUCGAUUACUAAAUG |
| M-044655-01 | D-044655-04 | Rad51c | 114714 | NM_053269 | CAGAAGGAGUCUACGAUAC |
| M-059082-01 | D-059082-01 | Rad51d | 19364 | NM_011235 | GCGCAGAUCUCUAUGAGGA |
| M-059082-01 | D-059082-02 | Rad51d | 19364 | NM_011235 | CAACGGGUCUGCAGGAGAU |
| M-059082-01 | D-059082-03 | Rad51d | 19364 | NM_011235 | GGAGCAGAGCCCAGAAUUA |
| M-059082-01 | D-059082-04 | Rad51d | 19364 | NM_011235 | GGAUACAGGUGGUGCGUUC |
| M-058519-00 | D-058519-01 | Xrcc2 | 57434 | NM_020570 | GAGCAGCCGGUUCUCAUUA |
| M-058519-00 | D-058519-02 | Xrcc2 | 57434 | NM_020570 | GCAGAAGCUCGUUGAAAGA |
| M-058519-00 | D-058519-03 | Xrcc2 | 57434 | NM_020570 | UUUAACAGCCCGAUGUAUA |
| M-058519-00 | D-058519-04 | Xrcc2 | 57434 | NM_020570 | GCGACAACGCAGAGUCUAA |
| M-045249-00 | D-045249-01 | Fbxo18 | 50755 | NM_015792 | GGAGGGAAAUAAUCAAUGA |
| M-045249-00 | D-045249-02 | Fbxo18 | 50755 | NM_015792 | AAACGAGACCUCAUCAUUA |
| M-045249-00 | D-045249-03 | Fbxo18 | 50755 | NM_015792 | GCAGAUCUAUACCUUCCGA |
| M-045249-00 | D-045249-04 | Fbxo18 | 50755 | NM_015792 | GGACUUUGCAGAGUACAUU |
| M-060606-01 | D-060606-01 | Chaf1a | 27221 | NM_013733 | GAGGAUGACUCCAUACUGA |
| M-060606-01 | D-060606-02 | Chaf1a | 27221 | NM_013733 | UCACACAGGCUCUCACGUA |
| M-060606-01 | D-060606-03 | Chaf1a | 27221 | NM_013733 | GAAAAGAGGGACCAGCAUA |
| M-060606-01 | D-060606-04 | Chaf1a | 27221 | NM_013733 | GCACGUGGGAUGUGUAUGG |
| M-061785-01 | D-061785-01 | Swsap1 | 66962 | XM_913186 | CCACAGCGCGAGAGCAUUG |
| M-061785-01 | D-061785-02 | Swsap1 | 66962 | XM_913186 | UGCCAUUACUUAAGCGAUA |
| M-061785-01 | D-061785-03 | Swsap1 | 66962 | XM_913186 | AGACGUCGCUGCUGUUUGC |
| M-061785-01 | D-061785-04 | Swsap1 | 66962 | XM_913186 | AGACAGGUGCAGAUUCAAA |
| M-060020-01 | D-060020-01 | Spidr | 224008 | NM_146068 | GCAAGAUGGUGUUUGGUUA |
| M-060020-01 | D-060020-02 | Spidr | 224008 | NM_146068 | CGAAAGAGAUCUGCCAUUU |
| M-060020-01 | D-060020-03 | Spidr | 224008 | NM_146068 | UAAGACACUUCCUGCAUUA |
| M-060020-01 | D-060020-04 | Spidr | 224008 | NM_146068 | GUUAUUGACUGGGAGGUUA |
| M-053866-01 | D-053866-01 | Fan1 | 330554 | XM_976717 | AGAGAUUGCCUCCGACUUA |
| M-053866-01 | D-053866-02 | Fan1 | 330554 | XM_976717 | AUUCAACUCUCGUCAGUAA |
| M-053866-01 | D-053866-04 | Fan1 | 330554 | XM_976717 | CGUAAAUUAACCUGGAUUA |
| M-053866-01 | D-053866-17 | Fan1 | 330554 | XM_976717 | CGAAAUUGUCCAGACGAUA |
| M-061987-01 | D-061987-01 | Blm | 12144 | NM_001042527 | GACACAAUCUGAAGUACUA |
| M-061987-01 | D-061987-02 | Blm | 12144 | NM_001042527 | CUAAAUCUAUGGAGGGUUA |
| M-061987-01 | D-061987-03 | Blm | 12144 | NM_001042527 | CCUAUGAUAUCGAUAACUU |
| M-061987-01 | D-061987-04 | Blm | 12144 | NM_001042527 | ACACCUGCGUUAAGUGAUA |
| M-064146-01 | D-064146-01 | Usp1 | 230484 | NM_146144 | GUUAUGAGCUUAUAUGUAG |
| M-064146-01 | D-064146-02 | Usp1 | 230484 | NM_146144 | CACAGUGGCAUUACUAUUA |
| M-064146-01 | D-064146-03 | Usp1 | 230484 | NM_146144 | CUACGACGAUGAAGUAUCA |
| M-064146-01 | D-064146-04 | Usp1 | 230484 | NM_146144 | AUUAUGAGCUGUACAACAA |
| M-059696-00 | D-059696-01 | Rmi1 | 74386 | NM_001168248 | GAAAGGACCCUCUUAAAUA |
| M-059696-00 | D-059696-02 | Rmi1 | 74386 | NM_001168248 | CAAACCAGCCCACGCAUUU |
| M-059696-00 | D-059696-03 | Rmi1 | 74386 | NM_001168248 | CUAGAAGGGUUACAGAAAU |

| | | | | | |
|---|---|---|---|---|---|
| M-059696-00 | D-059696-04 | Rmi1 | 74386 | NM_001168248 | GAAUGGAGUAUCAGUCUAU |
| M-052249-01 | D-052249-13 | Rmi2 | 223970 | NM_001162932 | GGAACUGGAUCCUCGGUUG |
| M-052249-01 | D-052249-14 | Rmi2 | 223970 | NM_001162932 | CUGCUUACAUGGACGCCUU |
| M-052249-01 | D-052249-15 | Rmi2 | 223970 | NM_001162932 | CCUGGUUCUUCGAAUGAUA |
| M-052249-01 | D-052249-16 | Rmi2 | 223970 | NM_001162932 | AGUAUGGCAUGGAUGUUAA |
| M-041565-01 | D-041565-02 | Slx1b | 75764 | NM_029420 | CGACCUGACUUCCGCCAUG |
| M-041565-01 | D-041565-03 | Slx1b | 75764 | NM_029420 | ACACAUGCCCAUUGCCUUU |
| M-041565-01 | D-041565-04 | Slx1b | 75764 | NM_029420 | UCGCAAGAAAGGUGGAGCA |
| M-041565-01 | D-041565-17 | Slx1b | 75764 | NM_029420 | GCGCACAUGCUUCGAGUUC |
| M-049483-00 | D-049483-01 | Smc1a | 24061 | NM_019710 | GUACAAGGGUCGACAGAUU |
| M-049483-00 | D-049483-02 | Smc1a | 24061 | NM_019710 | GAAAGGAGGCCAAACAAGA |
| M-049483-00 | D-049483-03 | Smc1a | 24061 | NM_019710 | GCAGGCAUUUGAACAGAUA |
| M-049483-00 | D-049483-04 | Smc1a | 24061 | NM_019710 | GAAAUUGGUGUGCGUAACA |
| M-055713-02-0005 | D-055713-14 | Rbbp8 | 225182 | NM_001081223 | CCUAGACACUGGCGUGAAA |
| M-055713-02-0005 | D-055713-15 | Rbbp8 | 225182 | NM_001081223 | GCAUUAACCGGCUACGAAA |
| M-055713-02-0005 | D-055713-16 | Rbbp8 | 225182 | NM_001081223 | AUAUUGAGGUAGUUCGGAA |
| M-055713-02-0005 | D-055713-17 | Rbbp8 | 225182 | NM_001081223 | AGAUAUGUUUGAUCGGACA |

**Supplementary Table S3:** Oligonucleotide list.

| Name | Purpose | Sequence (5' --> 3') |
|---|---|---|
| 1kb | sgRNA | TGTCGAGCCCGACGCGCGTG |
| mActinRTPCRP1 | primer (qPCR) | GGCTGTATTCCCCTCCATCG |
| mActinRTPCRP2 | primer (qPCR) | CCAGTTGGTAACAATGCCATGT |
| Top3aRTPCRUP | primer (qPCR) | TGGCTCATGACTTCCAGATG |
| Top3aRTPCRDN | primer (qPCR) | ATTGGGTTTTACAGCCTTGC |
| FancaRTPCRUP | primer (qPCR) | TGTCCCGTGATTCTGACTTC |
| FancaRTPCRDN | primer (qPCR) | ACTCCTCTCCACGCAAAGTG |
| Ep400RTPCRUP | primer (qPCR) | AAGGAAGGAAGGCTTGTGGT |
| Ep400RTPCRDN | primer (qPCR) | TCTTCCCTCTTTCCTCACGA |
| Msh6RTPCRUP | primer (qPCR) | AAGAAGCTGCCAGACCTTGA |
| Msh6RTPCRDN | primer (qPCR) | GCTTGAGGGTTTGGACGTA |
| Mlh1RTPCRUP | primer (qPCR) | GGTGGCTTCCTCATCCACTA |
| Mlh1RTPCRDN | primer (qPCR) | AGAGCAAGCATCTCCTCGTC |
| Pms2RTPCRUP | primer (qPCR) | ATAACGTGAGCTCCCCAGAA |
| Pms2RTPCRDN | primer (qPCR) | GAGGACCAGGCAATCTTTGA |
| Pms1RTPCRUP | primer (qPCR) | ATGGGCAACATGGAATCTGT |
| Pms1RTPCRDN | primer (qPCR) | TGGGATACAAGCGGGTAGAC |
| Smarcal1RTPCRUP | primer (qPCR) | AAATCCCATGTGTCGTCGAG |
| Smarcal1RTPCRDN | primer (qPCR) | GGCTGTGATGGACAACAGTG |
| FancmRTPCRUP3 | primer (qPCR) | CGAATCCTTTTCAGCTCTGG |
| FancmRTPCRDN3 | primer (qPCR) | AGGGGAGCTGTTAGCCATCT |
| Chaf1bRTPCRUP | primer (qPCR) | CTCCAATCTTGCTCGACACA |
| Chaf1bRTPCRDN | primer (qPCR) | CCCTCAGGGTCTTTACCACA |
| Zswim7RTPCRUP | primer (qPCR) | CAGGTGCTAGGGAGTTCTGG |
| Zswim7RTPCRDN | primer (qPCR) | GCCGCACCTTTTATCCTTCT |
| Ercc4RTPCRUP | primer (qPCR) | TGGAACAACACAAGCCTGAA |
| Ercc4RTPCRDN | primer (qPCR) | GCCACAAAGGGTCCAAGTAA |
| Topbp1RTPCRUP | primer (qPCR) | CACCTTGGAGCAAGTGTTCA |
| Topbp1RTPCRDN | primer (qPCR) | GTGCGTTGTCAACCAGAAAA |
| Ercc1RTPCRUP | primer (qPCR) | CCCTGAAAACAGGAGCAAAG |
| Ercc1RTPCRDN | primer (qPCR) | CACTTGAACCAGCAGCACAC |
| Pold3RTPCRUP | primer (qPCR) | GAGTGAGCGAAGCTGTTTCC |
| Pold3RTPCRDN | primer (qPCR) | CCCGAATTTTCTTTCCGTTT |
| Usp1RTPCRUP | primer (qPCR) | TGGATTATTTGCCGTTGTGA |
| Usp1RTPCRDN | primer (qPCR) | GGGCATTTCCACCAACTCTA |
| BlmRTPCRUP | primer (qPCR) | GGTCCAGAAGGACATCCTCA |
| BlmRTPCRDN | primer (qPCR) | CAGCCATTGTGTCACATTCC |
| CtIPRTPCRUP | primer (qPCR) | CCCAGGTACCAGATGAGGAA |
| CtIPRTPCRDN | primer (qPCR) | TACATGTGTGCCCAAGCAAT |
| Fancd2RTPCRUP3 | primer (qPCR) | TGATGAATTTGCCAACCTGA |
| Fancd2RTPCRDN3 | primer (qPCR) | GGCAGGAGGTTGATGACAAT |
| Xrcc3RTPCRUP3 | primer (qPCR) | CCCTGCTGAGACCACTTAGG |
| Xrcc3RTPCRDN3 | primer (qPCR) | GAAGCTCTCCTTCTGCTGGA |
| Slx4RTPCRUP | primer (qPCR) | CTTTATGCGAGATGCCCACT |
| Slx4RTPCRDN4 | primer (qPCR) | AGTCTCTGCTCGGCTTTCAC |
| Rmi1RTPCRUP1 | primer (qPCR) | TGGCTTTTGGGGTGTAACTG |
| Rmi1RTPCRDN1 | primer (qPCR) | AGCACCACCCCTTTACATGA |
| Rmi2RTPCRUP2 | primer (qPCR) | CTCCACCCCCTAGAAAAGC |
| Rmi2RTPCRDN2 | primer (qPCR) | AGCACCCCACGAACACATA |
| 1kbTIDEUP1 | TIDE | gctcgtagaaggggaggttg |
| 1kbTIDEDN1 | TIDE | atagcagctttgctccttcg |

| | | |
|---|---|---|
| 1kbTIDEUP2 | TIDE | caggaggccttccatctgt |
| 1kbTIDEDN2 | TIDE | GGCCCCATTATTGAAGCATT |
| 16bpTIDEUP1 | TIDE | attaagggccagctcattcc |
| 16bpTIDEDN1 | TIDE | TGGTGCAGATGAACTTCAGG |
| 9kbTIDEUP1 | TIDE | CCATGGGCCAGTAGAATGAC |
| 9kbTIDEDN1 | TIDE | TTTGAGAAGGTGGCTGTCCT |
| 9kbTIDEUP2 | TIDE | AGCATCCCTAACCTGGAAGC |
| 9kbTIDEDN2 | TIDE | AAAGATCTGCTTGCCTCTGC |
| RMDjunct368UPillumina | primer for strand bias | ACACTCTTTCCCTACACGACGCTCTTCCGATCTCCGGGTCCTTCTTGTGTTTC |
| RMDjunct368DNillumina | primer for strand bias | GACTGGAGTTCAGACGTGTGCTCTTCCGATCTAACAGCTCCTCGCCCTTG |
| Mlh1e11sgAUP | sgRNA | caccgCATTGACGTCCACGTTCTGA |
| Mlh1e11sgBUP | sgRNA | caccgCGAAGTTCACTTTCTGCACG |

**Supplemental Figure S1: Depletion of BLM and CtIP via siRNA. (A)** Immunoblotting analysis of BLM, CTIP, and ACTIN in WT mESCs transfected with either siCTRL, siBlm, or siCtIP siRNAs. **(B)** Shown is qRT-PCR analysis of BLM and CtIP in WT mESCs transfected with either siCTRL, siBlm, or siCtIP siRNAs. Shown is the mRNA abundance of BLM and CtIP based on a threshold cycle (Ct) values from PCR amplification, normalized to ACTIN, relative to siCTRL treated cells (siCTRL = 1). n=3 PCR. $^{**}p \leq 0.005$, $^{***}p \leq 0.0005$, unpaired t-test. Data are represented as mean values ± SD.

**Supplemental Figure S2: qRT-PCR analysis of several siRNAs, and reporter analysis of 3 genes that did not show significant effects on RMDs vs. NHEJ. (A)** qRT-PCR analysis of siRNAs targeting the 18 genes described in Figure 2. From siRNA treatment in WT mESCs, shown is the mRNA abundance of each gene based on a threshold cycle (Ct) values from PCR amplification, normalized to actin, relative to siCTRL treated cells (siCTRL = 1). n=3 PCR. $^{*}p \leq 0.05$, $^{**}p \leq 0.005$, $^{***}p \leq 0.0005$, $^{****}p < 0.0001$, Statistics unpaired t-test. **(B)** Effects of siRNAs targeting Ep400, Zswim7, and Ercc1 on the four RMD events and NHEJ. Frequencies are normalized to transfection efficiency and parallel siCTRL (=1). n=4. $^{*}p \leq 0.05$, one-way ANOVA using Tukey's multiple comparisons test. Also shown is qRT-PCR analysis of siRNA treatments targeting these genes, as in (A). $^{**}p \leq 0.005$, $^{***}p \leq 0.0005$, Statistics unpaired t-test, n=3 PCR. Data are represented as mean values ± SD.

**Supplemental Figure S3: Frequency of indels with the sgRNAs targeting the 16 bp, 1 kbp, and 9.1 kbp DSBs.** Shown is Tracking of Indels by DEcomposition (TIDE) analysis of samples following single DSB by sgRNAs targeting the 16 bp, 1 kbp, or 9.1 kbp DSB sites. n=3. Data are represented as mean values ± SD.

**Supplemental Figure S4: MLH1 suppresses RMDs independent of DSB/repeat distance, and validation of MLH1 and MSH6 siRNAs. (A)** The RMD frequencies for *Mlh1*$^{-/-}$ shown in Figure 2C were normalized to WT (=1), and grouped by sequence divergence between the repeats to enable comparisons across DSB/repeat distances. n=6. *$p \leq 0.05$, **$p \leq 0.005$, ***$p \leq 0.0005$, ****$p < 0.0001$, one-way ANOVA with Tukey's multiple comparisons test. Data are represented as mean values ± SD. **(B)** Immunoblotting analysis of MLH1 and ACTIN in WT, *Msh2*$^{-/-}$, *Mlh1*$^{-/-}$, and *Exo1*$^{-/-}$ mESCs that were treated with siMlh1 and siCTRL. **(C)** Immunoblotting analysis of MSH6 and ACTIN in WT, *Msh2*$^{-/-}$, *Mlh1*$^{-/-}$, and *Exo1*$^{-/-}$ mESCs that were treated with siMsh6 and siCTRL.

**Supplemental Figure S5: Suppression of RMDs via PMS2 and PMS1 is MLH1-dependent. (A)** Shown is the effects of loss of PMS2 and PMS1 individually, and in combination, on RMD events between identical repeats with the 9.1 kbp and 16 bp DSB/repeat distances. Frequencies are normalized to transfection efficiency and parallel siCTRL (=1). n = 6. ***$p \leq 0.0005$, siCTRL vs. each set of siRNAs, and the combination siRNA treatment (siPms2 + siPms1) vs. the individual gene siRNAs, using unpaired t-tests with Holm-Sidak correction. **(B)** Shown is the effect of treatment of siRNAs targeting both PMS2 and PMS1 (siPms1 + siPms2) in *Mlh1*$^{-/-}$ mESCs on the three RMD reporters, each tested with the 9.1 kbp and 16 bp DSB/repeat distances. Frequencies are normalized to transfection efficiency and parallel siCTRL (=1). n=6. **$p \leq 0.005$, unpaired t-test. **(C)** Shown is qRT-PCR analysis of PMS2 and PMS1 in WT mESCs transfected with either siCTRL, siPms2, siPms1, or both siPms2 + siPms1. Shown is the mRNA abundance of Pms1 and Pms2, based on a threshold cycle (Ct) values from PCR amplification, normalized to actin, relative to siCtrl treated cells (siCTRL = 1). n=3 PCR. *$p \leq 0.05$, **$p \leq 0.005$, unpaired t-test. **(D)** Shown is qRT-PCR analysis of PMS2 and PMS1 in *Mlh1*$^{-/-}$ mESCs upon transfection with siRNAs targeting PMS2 and PMS1 in combination. Analysis as in (C). Notably, while siRNA targeting PMS2 only affected PMS2 RNA levels, siRNA targeting PMS1 caused a modest but consistent decrease in PMS2 RNA. Although, the combination of siRNA targeting PMS1 and PMS2 showed a similar depletion of PMS2 RNA as the siRNAs targeting PMS2 alone. In contrast, as described in the Results, the combination siRNA targeting of PMS1 and PMS2 caused the greatest increase in RMDs. n=3. **p ≤ 0.005, ***p ≤ 0.0005, Statistics are unpaired t-test. Data are represented as mean values ± SD.

**Supplemental Figure S6: Effect of TOP3α Y362F on RMD frequency. (A)** Shown are effects of expressing TOP3α WT and Y362F in cells treated with siTop3 for 6 RMD events: RMD-GFP, 1%RMD-GFP, and 3%RMD-GFP, each with the 9.1 kbp and 16 bp DSB/repeat distances. Frequencies are normalized to transfection efficiency and parallel siCTRL (=1). n=6. *p ≤ 0.05, **p ≤ 0.005, ***p ≤ 0.0005, ****p < 0.0001, siCTRL EV vs. siTop3a EV unpaired t-test, and siTop3a EV vs. siTop3a TOP3α and siTop3a TOP3α vs. siTop3a TOP3α-Y362F unpaired t-tests with Holm-Sidak correction. Data are represented as mean values ± SD. **(B)** Immunoblotting analysis of TOP3A and ACTIN in WT mESCs transfected with either siCtrl EV, or siTop3a with EV, WT TOP3α, or Y362F. Same blot as in Figure 3D but with un-cropping the Y362F lane.

**Supplemental Figure S7**: **Influence of sequence divergence and DSB/repeat distance on the relative fold effect of TOP3α on RMDs, and assessment of TOP3α depletion in various contexts. (A)** RMD frequencies from siTop3α treated cells shown in Figure 6A were normalized to siCTRL (=1), and grouped by sequence divergence to enable comparisons of effects of DSB/repeat distance. n=6. **p ≤ 0.005, ***p ≤ 0.0005, ****p < 0.0001, one-way ANOVA with Tukey's multiple comparisons test. **(B)** Analysis as in (A) except grouped by DSB/repeat distances to enable comparisons with varying degrees of sequence divergence. n=6. *p ≤ 0.05, **p ≤ 0.005, ***p ≤ 0.0005, ****p < 0.0001, one-way ANOVA with Tukey's multiple comparisons test. **(C)** Shown is qRT-PCR analysis of TOP3α in WT, *Mlh1*$^{-/-}$, and *Msh2*$^{-/-}$ mESCs upon transfection with siRNAs targeting TOP3α. Shown is the mRNA abundance of TOP3α based on a threshold cycle (Ct) values from PCR amplification, normalized to actin, relative to siCTRL treated cells (siCTRL = 1). n=3 PCR. ***p ≤ 0.0005, unpaired t-test.

Suppl Figure S1 Trost et al.

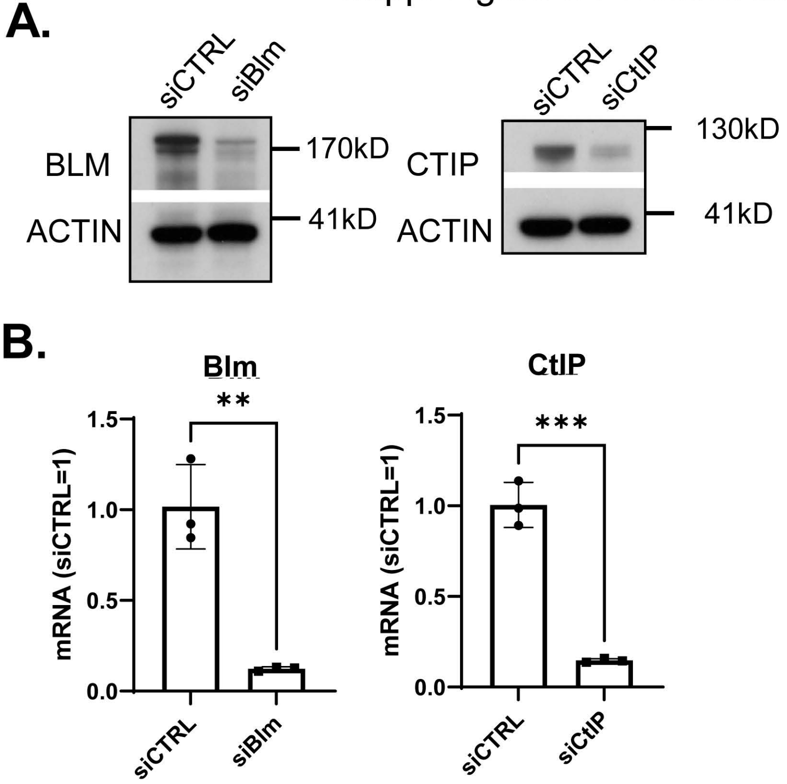

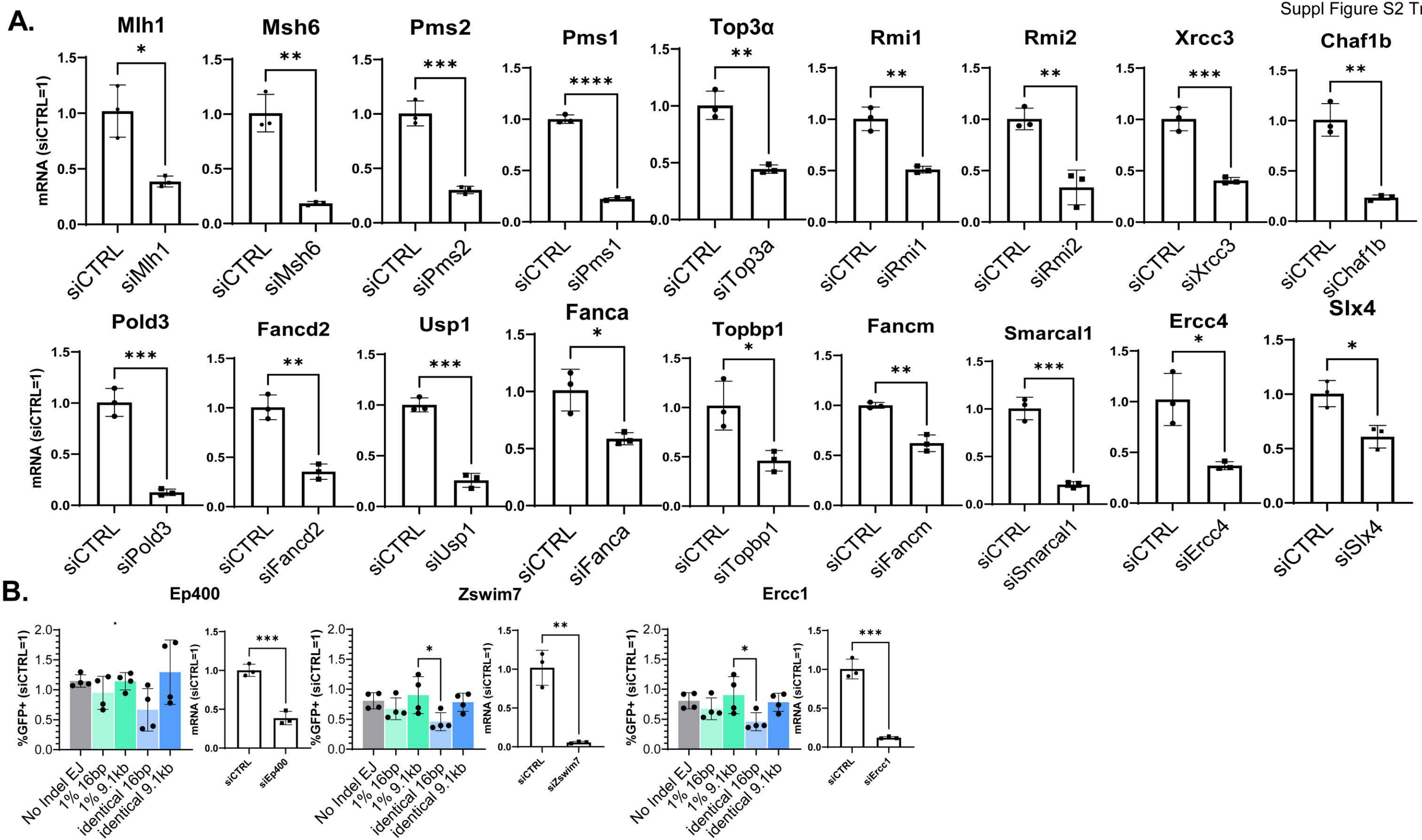
A.
Mlh1
Msh6
Pms2
Pms1
Top3α
Rmi1
Rmi2
Xrcc3
Chaf1b
mRNA (siCTRL=1)
siCTRL
siMlh1
siMsh6
siPms2
siPms1
siTop3a
siRmi1
siRmi2
siXrcc3
siChaf1b
Pold3
Fancd2
Usp1
Fanca
Topbp1
Fancm
Smarcal1
Ercc4
Slx4
siPold3
siFancd2
siUsp1
siFanca
siTopbp1
siFancm
siSmarcal1
siErcc4
siSlx4
B.
Ep400
Zswim7
Ercc1
%GFP+ (siCTRL=1)
No Indel EJ
1% 16bp
1% 9.1kb
identical 16bp
identical 9.1kb
siEp400
siZswim7
siErcc1

Suppl Figure S3 Trost et al

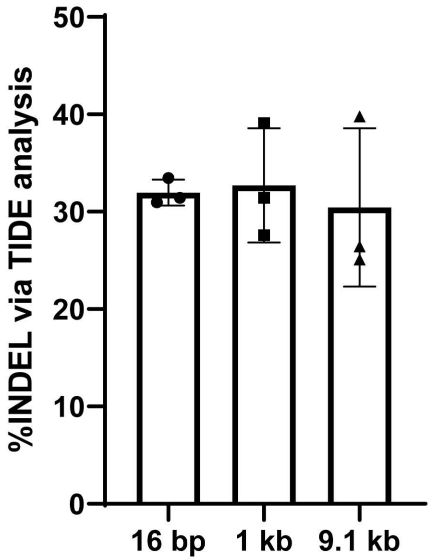

Suppl Figure S4 Trost et al.

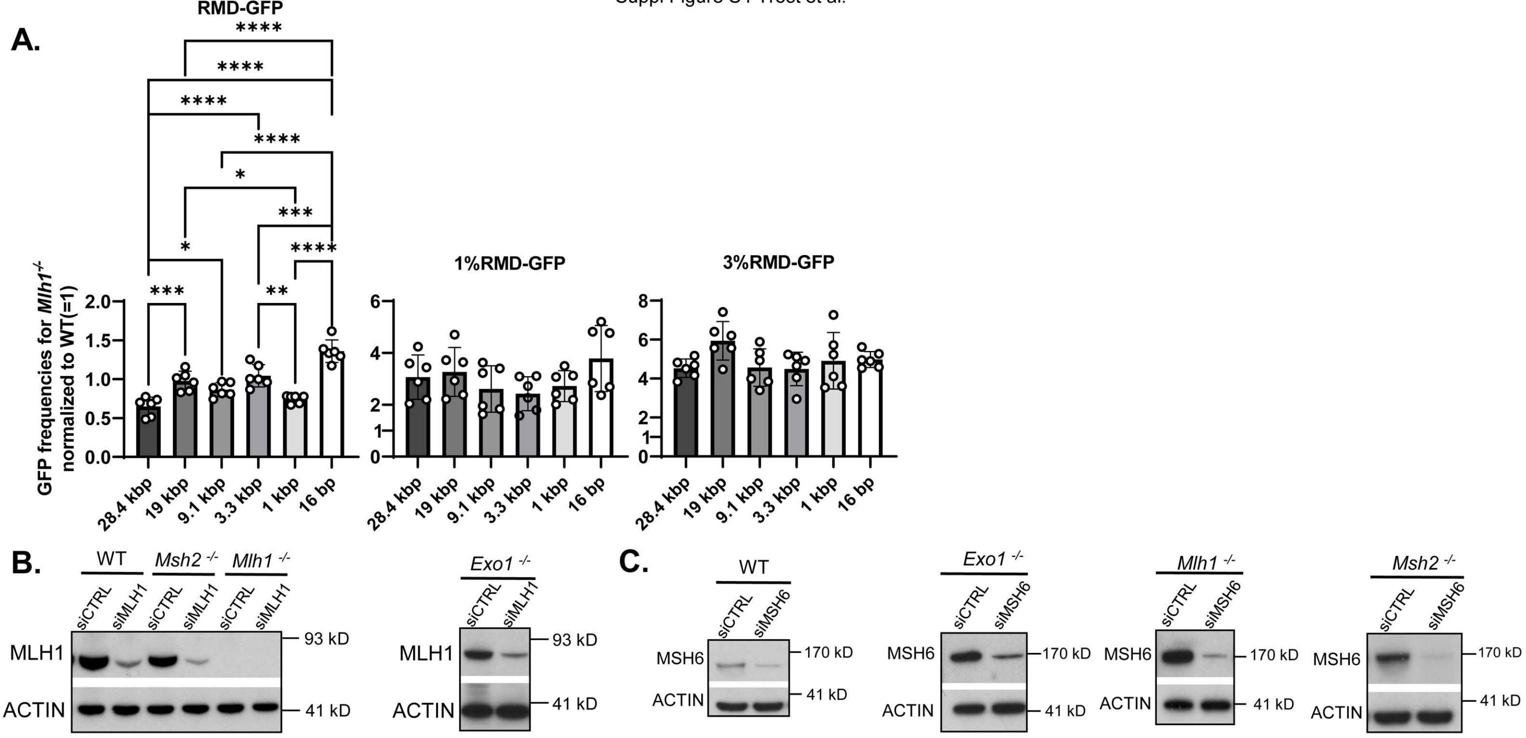

Suppl Figure S5 Trost et al.

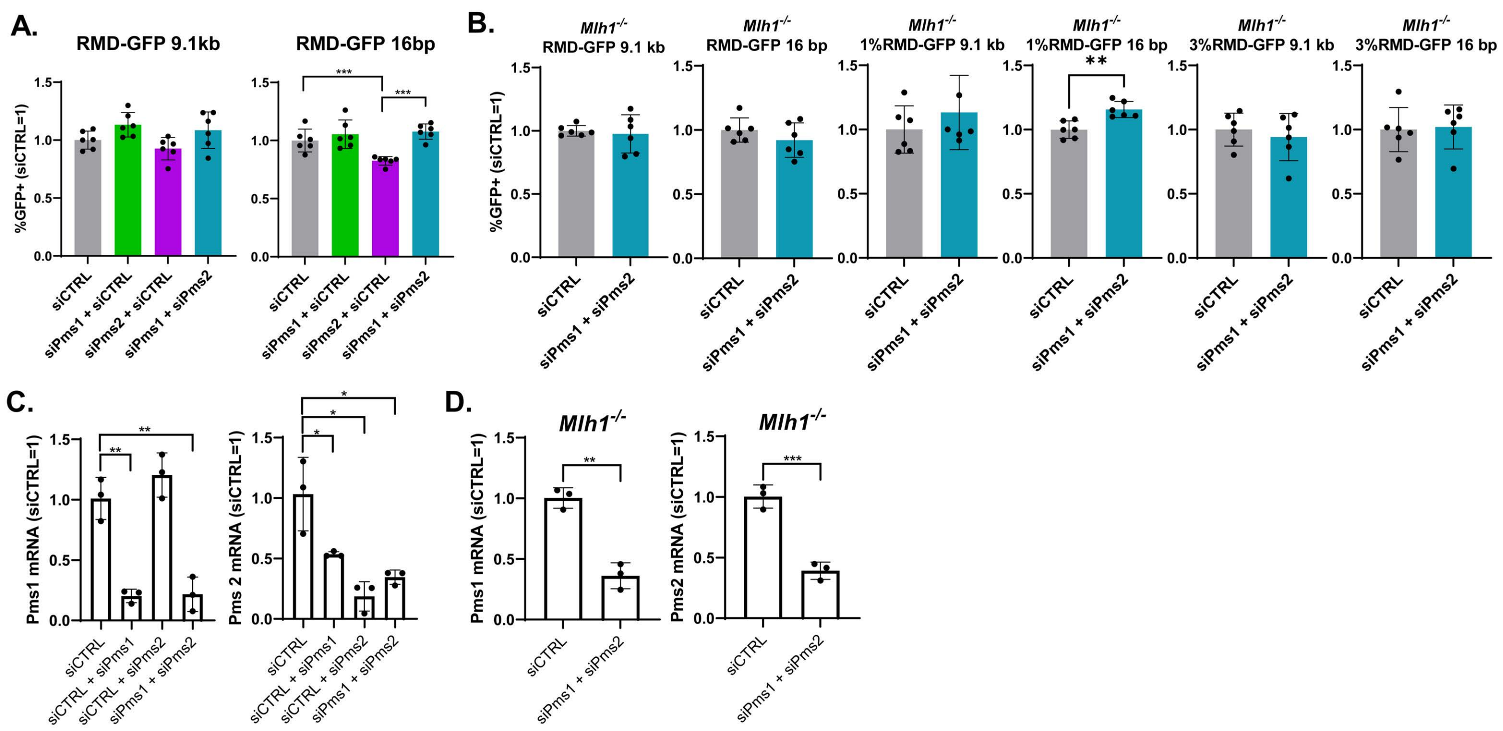

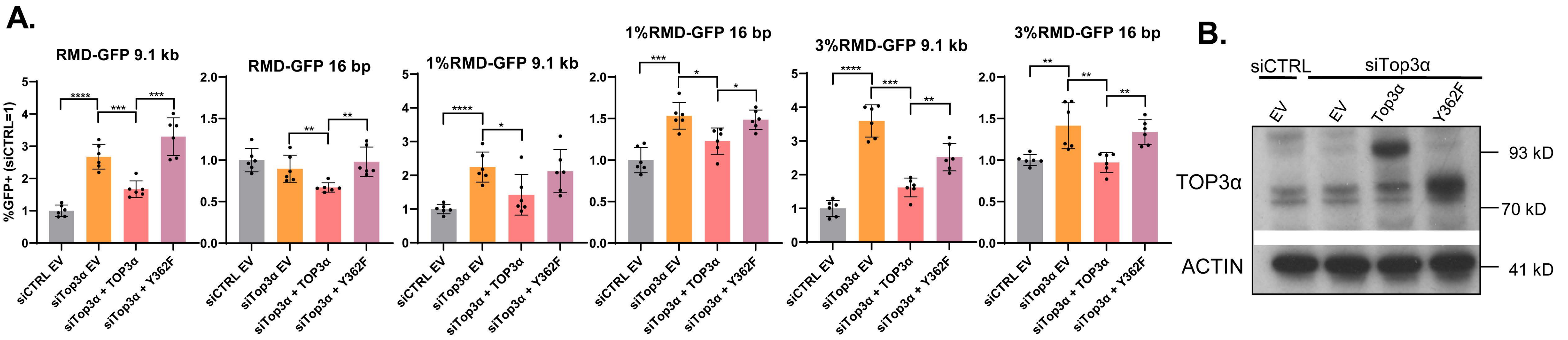
A.
RMD-GFP 9.1 kb
RMD-GFP 16 bp
1%RMD-GFP 9.1 kb
1%RMD-GFP 16 bp
3%RMD-GFP 9.1 kb
3%RMD-GFP 16 bp
%GFP+ (siCTRL=1)
siCTRL EV
siTop3α EV
siTop3α + TOP3α
siTop3α + Y362F
B.
siCTRL
siTop3α
EV
EV
Top3α
Y362F
TOP3α
ACTIN
93 kD
70 kD
41 kD

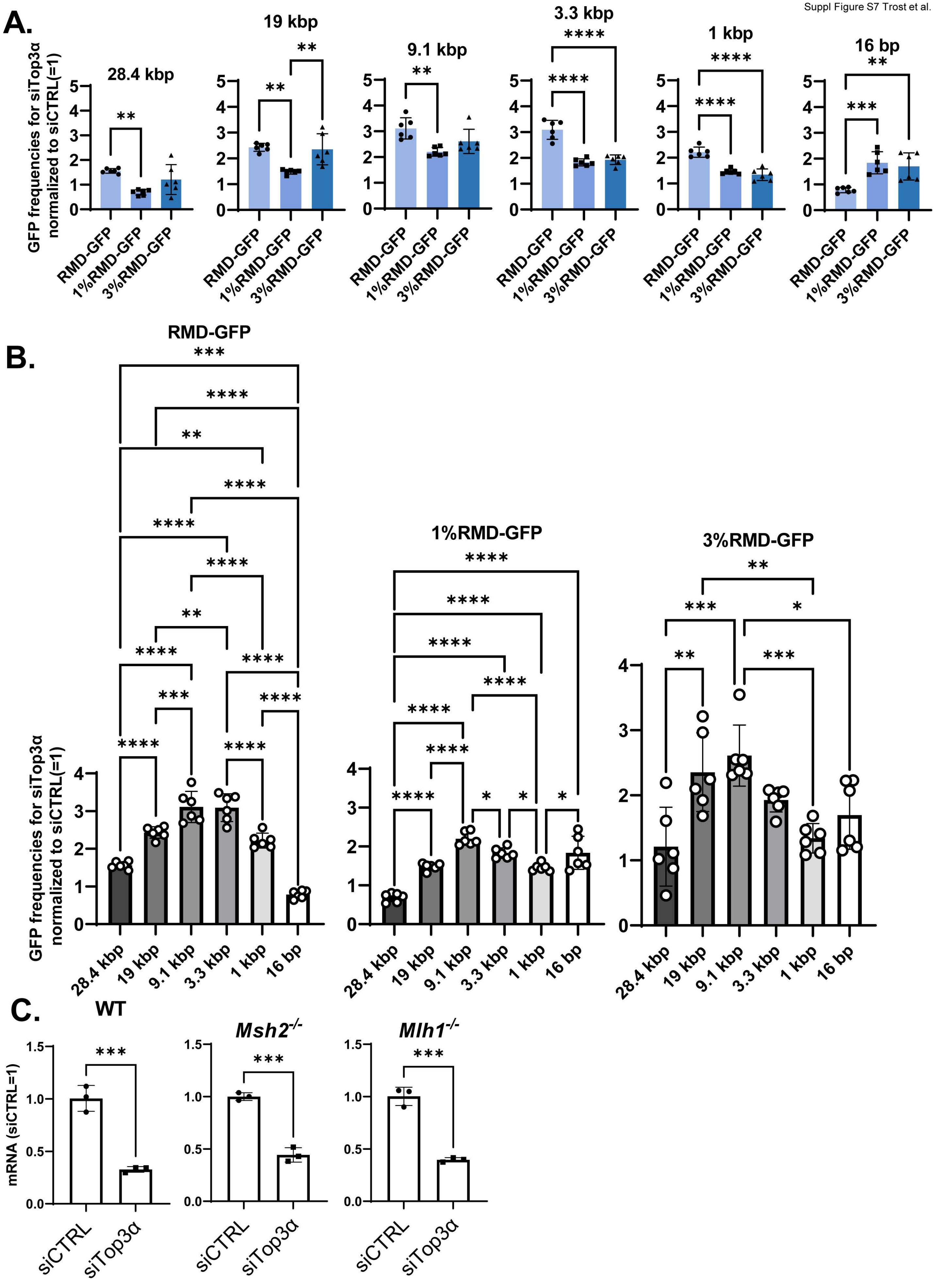
A.
28.4 kbp
19 kbp
9.1 kbp
3.3 kbp
1 kbp
16 bp
GFP frequencies for siTop3α normalized to siCTRL(=1)
RMD-GFP
1%RMD-GFP
3%RMD-GFP
B.
RMD-GFP
1%RMD-GFP
3%RMD-GFP
GFP frequencies for siTop3α normalized to siCTRL(=1)
28.4 kbp
19 kbp
9.1 kbp
3.3 kbp
1 kbp
16 bp
C.
WT
Msh2-/-
Mlh1-/-
mRNA (siCTRL=1)
siCTRL
siTop3α
